# Myonuclear domain-associated and central nucleation-dependent spatial restriction of dystrophin protein expression in a novel DMD mouse model

**DOI:** 10.1101/2025.07.24.666562

**Authors:** Katarzyna Chwalenia, Vivi-Yun Feng, Nicole Hemmer, John C.W. Hildyard, Liberty E. Roskrow, Richard J. Piercy, Eric T. Wang, Annemieke Aartsma-Rus, Maaike van Putten, Matthew J.A. Wood, Thomas C. Roberts

## Abstract

The restoration of uniformly-distributed dystrophin protein expression is an important consideration for the development of advanced therapeutics for Duchenne muscular dystrophy (DMD). To explore this concept, we generated a novel genetic mouse model (*mdx52-Xist*^Δhs^) that expresses variable, and non-uniformly distributed, dystrophin protein from birth as a consequence of skewed X-chromosome inactivation. *mdx52-Xist*^Δhs^ myofibers are heterokaryons containing a mixture of myonuclei expressing either wild-type or mutant dystrophin alleles in a mutually exclusive manner, resulting in dystrophin protein being spatially restricted to corresponding dystrophin-expressing myonuclear domains. This phenotype models the situation in female DMD carriers, and dystrophic muscle in which dystrophin has been incompletely restored by partially-effective experimental therapeutics. Total dystrophin expression increased in aged (60-week-old) *mdx52-Xist*^Δhs^ mice relative to 6-week-old adults, suggestive of an accumulation of dystrophin-expressing myonuclei through positive selection, although this was insufficient to resolve sarcolemmal dystrophin patchiness. Nevertheless, compared to mice expressing no dystrophin, non-uniformly-distributed dystrophin was protective against pathology-related muscle turnover in an expression-level-dependent manner in both adult and aged *mdx52-Xist*^Δhs^ mice. Systematic classification of isolated *mdx52-Xist*^Δhs^ myofibers revealed profound differences associated with central nucleation, with dystrophin found to be translationally repressed in centrally-nucleated myofibers and myofiber segments. These findings have important implications for the development of dystrophin restoration therapies.

## Introduction

DMD is a monogenic muscle wasting disorder caused by pathogenic variants (frequently whole exon deletions) in the *DMD* gene, which encodes the dystrophin protein. Dystrophin forms a mechanical link between the cytoskeleton (i.e. filamentous actin and microtubules) and the extracellular matrix via interactions with components of the dystrophin-associated protein complex (DAPC), which consists of both structural and signalling proteins. Absence of dystrophin protein at the sarcolemma, and subsequent disruption of DAPC assembly, sensitizes muscle to contraction-induced damage.^1^ In DMD patients, this leads to perpetual muscle turnover, persistent inflammation, and the progressive replacement of myocytes with fatty and fibrotic tissue.

While still rare, the relatively high prevalence of the disorder (~1 in 5,000 males) and the severity of the disease have made DMD a priority candidate for experimental therapeutics. Indeed, there are now four antisense oligonucleotide (ASO) drugs and one microdystrophin gene therapy that have received (accelerated) marketing authorization from the US FDA.^2^ Furthermore, a multitude of other approaches are under investigation, including CRISPR-Cas9-mediated gene editing and upregulation of utrophin (a dystrophin paralogue).^2–7^ Despite this progress, the clinical challenge of effectively treating DMD remains incompletely met, in part due to a combination of poor drug delivery, incomplete functionality of the restored internally-deleted quasi-dystrophin protein, and failure to rescue dystrophin in all fibers. We have observed that the pattern of dystrophin expression restored following treatment is dependent on the modality. Specifically, we observed a uniform sarcolemmal pattern of dystrophin expression following treatment of the *mdx* mouse model of DMD with peptide-phosphorodiamidate morpholino oligonucleotides (PPMOs) designed to induce exon skipping of *Dmd* exon 23 (containing a premature termination codon),^8,9^ and a patchy pattern of dystrophin expression following CRISPR-Cas9-mediated excision of the same exon in the severely-affected dystrophin/utrophin double knock-out (dKO) mouse.^10^ Similar results were also reported by Morin *et al*.^11^ Importantly, analysis of human biopsies has shown that incomplete sarcolemmal dystrophin coverage correlates with pathological severity in Becker muscular dystrophy and intermediate muscular dystrophy patients (i.e. those with clinical phenotypes that are between those of BMD and DMD).^12^ Spatial restriction of dystrophin can be attributed to its limited capacity to diffuse throughout the sarcolemma, such that it becomes localized in the vicinity of its corresponding myonucleus of origin, consistent with the myonuclear domain hypothesis.^13^

We have previously modelled the effects of patchy sarcolemmal dystrophin expression using skewed X-chromosome inactivation (XCI) in a murine system.^9,14,15^ To further investigate this patchy dystrophin phenomenon, we have developed a novel mouse model that exhibits preferential XCI of the healthy X-chromosome, while the mutated X-chromosome carries a patient-relevant whole exon deletion of *Dmd* exon 52.^16^ Using this novel system, we show that the female *mdx52-Xist*^Δhs^ mice exhibit variable levels of dystrophin expression with a characteristic patchy pattern of dystrophin coverage at the sarcolemma. Comparison of *mdx52-Xist*^Δhs^ mice at adult (6 week) and aged (60 week) time points revealed an overall increase in total dystrophin level, consistent with the accumulation of dystrophin-positive myofibers with time. However, this increase in dystrophin expression was insufficient to resolve sarcolemmal dystrophin patchiness, suggesting that these fibers are incompletely protected from the cycles of myonecrosis and compensatory regeneration that are characteristic pathological features of DMD. However, myofiber central nucleation was inversely correlated with total dystrophin expression, suggesting that patchy dystrophin expression does offer myofibers a degree of protection. Interestingly, analysis of *mdx52-Xist*^Δhs^ isolated single myofibers revealed that centrally-nucleated myofibers and myofiber segments were almost completely devoid of dystrophin or DAPC expression. This unexpected finding was attributed to local and specific inhibition of dystrophin expression at the level of translation. This study has important implications for therapeutic efforts to restore dystrophin protein expression in Duchenne patients.

## Results

### Dystrophin is expressed in a within-fiber patchy manner in adult *mdx52-Xist*^Δhs^ muscle

To generate a genetic mouse model with patchy sarcolemmal dystrophin protein expression we bred male *mdx52* mice^16,17^ (which carry a patient-relevant deletion of *Dmd* exon 52, leading to disruption of dystrophin expression) and female *Xist*^Δhs^ mice^18^ (which carry a deletion in a DNase I hypersensitivity site within the *Xist* promoter, leading to skewed XCI of the host chromosome). The resulting female F1 progeny (*mdx52-Xist*^Δhs^) are expected to express dystrophin at variable levels as a consequence of skewed (i.e. preferential) XCI of the healthy *Dmd* allele (**Figure S1**).^9,14,15^ This mouse is a model of (i) female dystrophinopathy (previously known as manifesting carriers),^19,20^ and (ii) the situation in dystrophic muscle following partial CRISPR-Cas9 correction.^9,10^

Analysis of *mdx52-Xist*^Δhs^ females (*N*=20) revealed a range of total dystrophin expression levels in 6-week-old tibialis anterior (TA) muscles and mice were retrospectively assigned to high (~23-41% of WT dystrophin levels, *n*=4), medium (~11-17%, *n*=7), and low (~1-8%, *n*=9) dystrophin-expressing groups *post-mortem*, as determined by western blot (**Figure 1A-C**). The distributions of dystrophin expression were consistent with those reported in similar studies by our groups.^9,14^ Immunofluorescence analysis in the same tissues revealed a within-myofiber patchy pattern of sarcolemmal dystrophin (**Figure 1D**). Regions of adjacent dystrophin-positive and dystrophin-negative sarcolemma were observed in the *mdx52*-*Xist*^Δhs^ muscles at all dystrophin expression levels. By contrast, dystrophin was uniformly-distributed in age- and sex-matched wild-type C57 and *Xist*^Δhs^ mice, and absent in *mdx52* controls (**Figure 1D**). Patchiness was most apparent in longitudinal sections, but incomplete sarcolemmal coverage was also apparent in some myofibers in transverse sections (especially in the low dystrophin *mdx52-Xist*^Δhs^ group). These data show that dystrophin mRNA and protein are not free to diffuse freely within syncytial myofibers, consistent with previous reports.^9–11^

**Figure 1.**
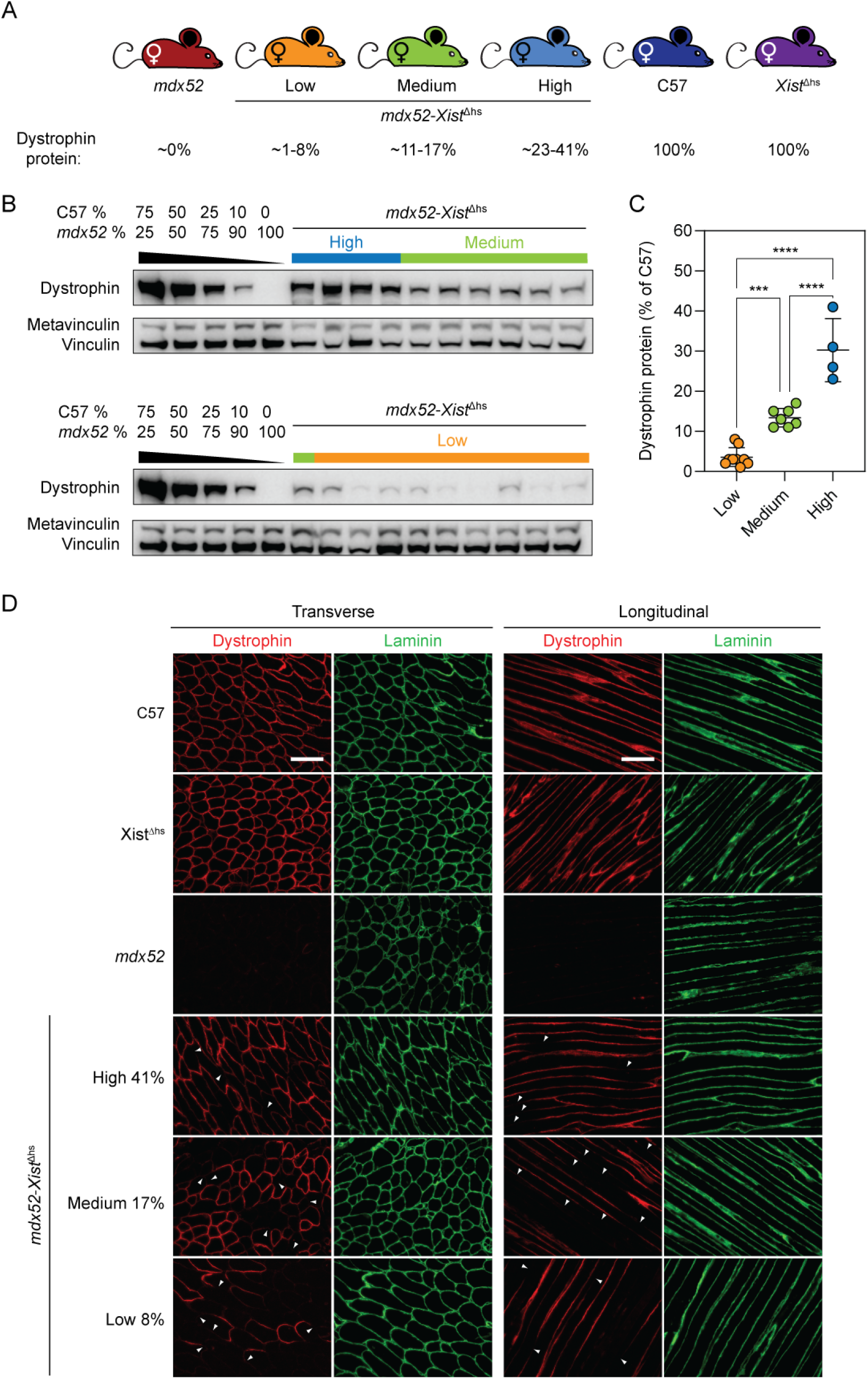
Dystrophin expression is patchy in *mdx52-Xist*^Δhs^ muscle. (**A**) *mdx52-Xist*^Δhs^ animals were assigned to high (*n*=4), medium (*n*=7), and low (*n*=9) dystrophin expressing groups post-mortem based on protein quantification in 6-week-old tibialis anterior (TA) muscles as determined by (**B**) Western blot analysis. Wild-type C57and *Xist*^Δhs^ mice were used as 100% dystrophin-expressing controls, and *mdx52* mice used as ~0% dystrophin-expressing controls. Vinculin was used as a loading control. (**C**) Dystrophin was quantified by comparison to standard curves containing defined mixtures of C57 and *mdx52* TA lysates. (**D**) Representative immunofluorescence staining of dystrophin and laminin in transverse and longitudinal TA muscle sections of 6-week-old C57 wild-type, *Xist*^Δhs^, *mdx52*, and *mdx52-Xist*^Δhs^ animals from high, medium, and low dystrophin-expressing groups. Within-fiber, patchy dystrophin expression resulting from skewed X-chromosome inactivation indicated with arrowheads. Scale bars indicate 100 µm, images taken at 20× magnification. The percentage values indicate total dystrophin quantification in the animals from which the sections were derived. Values are mean+SD. Statistical significance was assessed by one-way ANOVA with Bonferroni *post hoc* test, *** *P*<0.001, \*\*\*\**P*<0.0001.

### Dystrophin, β-dystroglycan, and α-dystrobrevin are localized in sarcolemmal patches in *mdx52-Xist*^Δhs^ isolated single myofibers

Analysis of *mdx52-Xist*^Δhs^ isolated single extensor digitorum longus (EDL) myofibers revealed similar patchy sarcolemmal distributions for dystrophin and the DAPC components β-dystroglycan (DAG1) and α-dystrobrevin (DTNA) (**Figure 2A**). Dual staining showed that dystrophin and β-dystroglycan were co-localized to common regions of the sarcolemma, forming a ‘zebra-like’ banding pattern of staining (**Figure 2B**). Conversely, the DAPC protein neuronal nitric oxide synthase (nNOS, NOS1) was uniformly distributed throughout single isolated myofibers derived from both *mdx52-Xist*^Δhs^ and WT C57 controls (**Figure 2C**).

**Figure 2.**
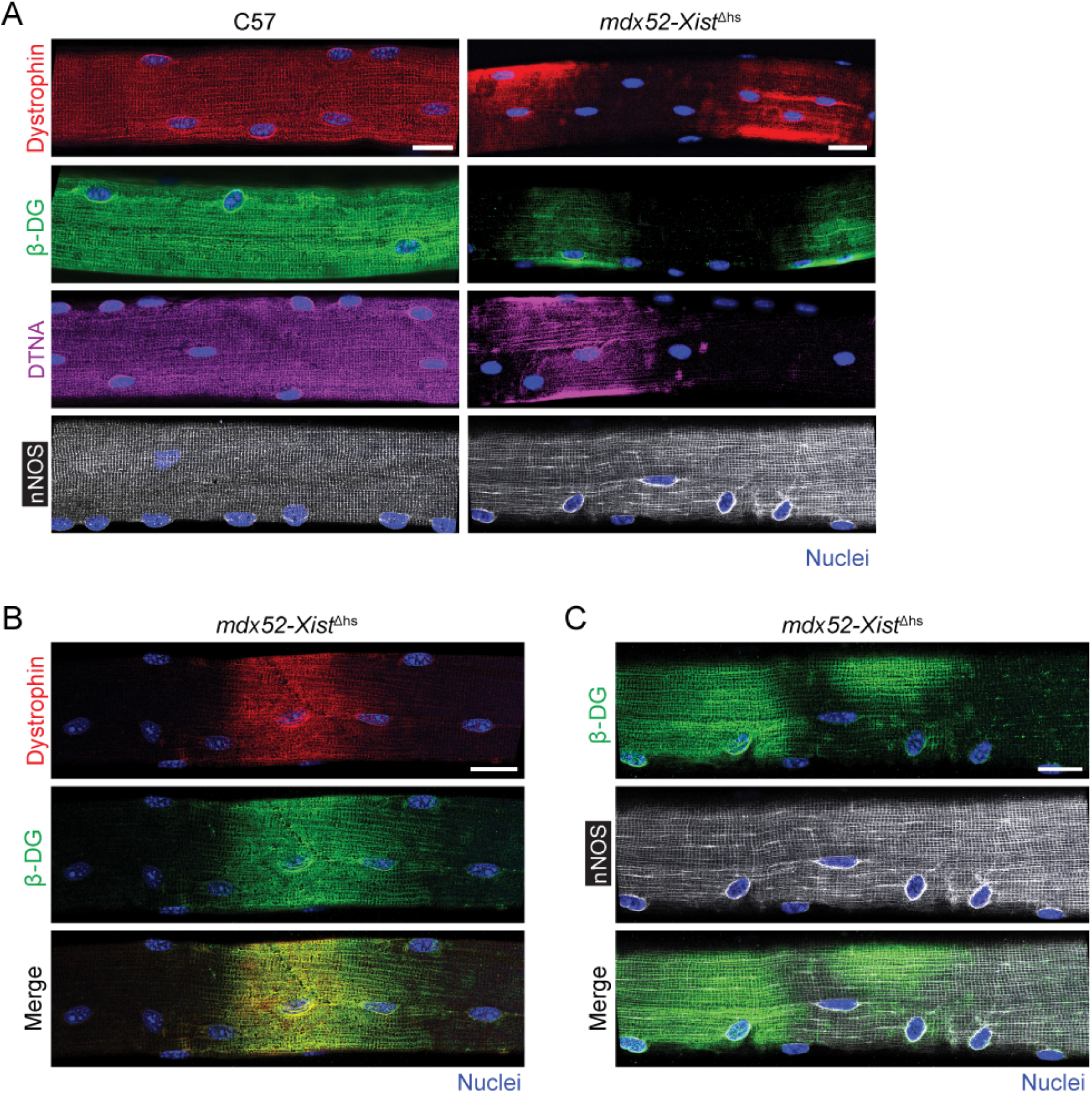
Dystrophin and DAPC protein expression is patchy in *mdx52-Xist*^Δhs^ isolated single myofibers. (**A**) Representative immunofluorescence staining of β-dystroglycan (β-DG), α-dystrobrevin (DTNA) and neuronal nitric oxide synthase (nNOS) in single isolated extensor digitorum longus (EDL) myofibers of adult (6-12 weeks old) C57 (wild-type) and *mdx52-Xist*^Δhs^ mice. Representative co-staining images of (**B**) dystrophin and β-DG, and (**C**) β-DG and nNOS in the isolated single myofiber of *mdx52-Xist*^Δhs^ animals. Scale bars indicates 20 µm, images taken at 25× magnification. Nuclei were stained with DAPI.

### Dystrophin expression is inversely correlated with muscle histopathology in adult *mdx52-Xist*^Δhs^ muscles

Histopathological analysis in adult *mdx52-Xist*^Δhs^ TA muscles revealed the presence of abundant centrally-nucleated fibers (CNFs) and foci of small diameter regenerating fibers (**Figure 3A**) at all levels of dystrophin expression, indicative of ongoing or historic muscle turnover. Mean CNF values were 11.2%, 16.7%, and 38.7% for high, medium, and low dystrophin expressing muscles, respectively (**Figure 3B**). The percentage of CNFs was strongly inversely correlated with dystrophin expression (Spearman’s *r*=-0.86, *P*=0.0023, **Figure 3C**). Myofiber size distributions were similar between all analysed genotypes (**Figure 3D**).

**Figure 3.**
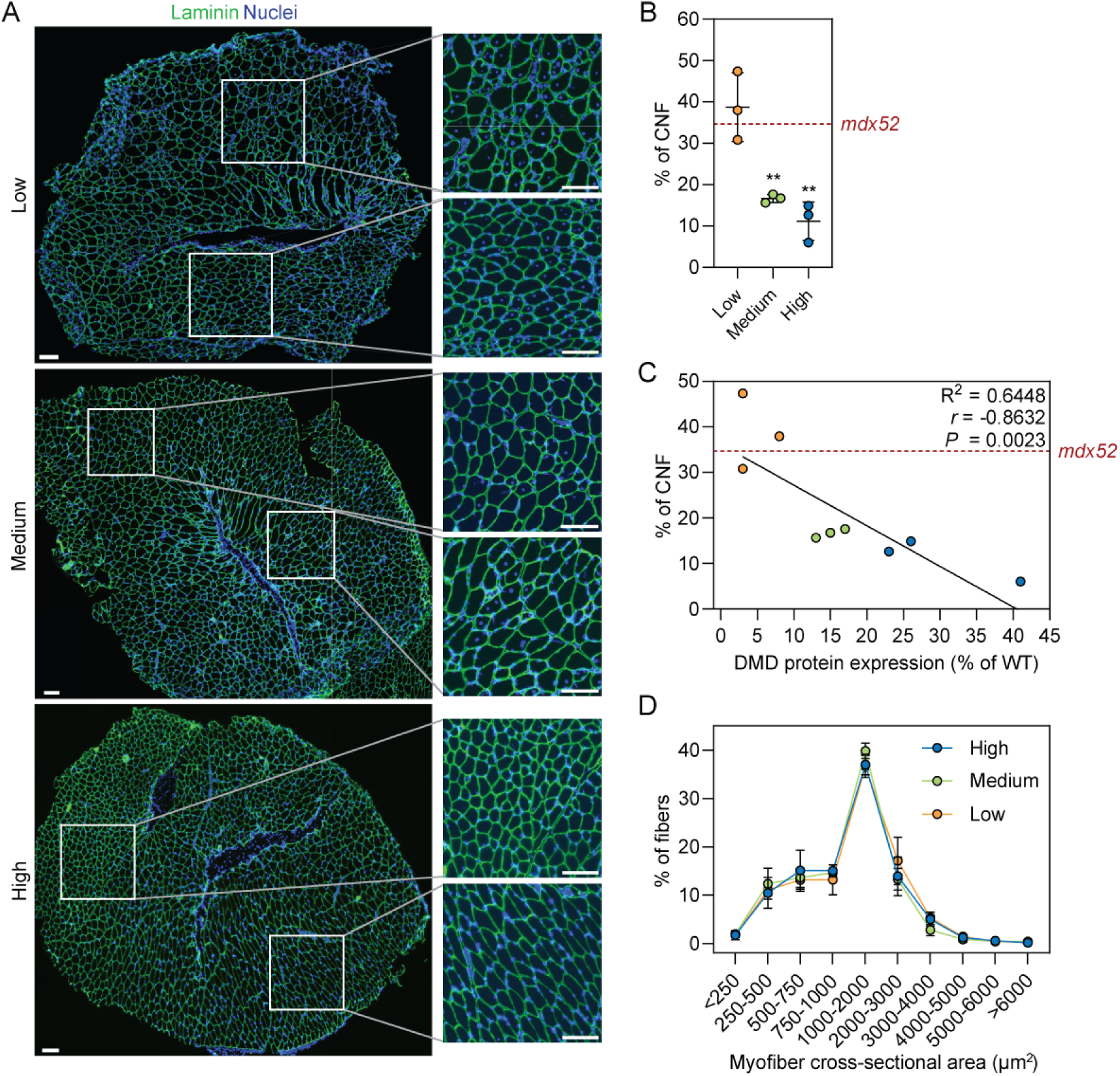
Dystrophin expression is inversely correlated with histopathology in *mdx52-Xist*^Δhs^ muscle sections. (**A**) Representative immunofluorescence images of transverse TA muscle sections from adult (6-week-old) *mdx52-Xist*^Δhs^ stained for laminin (green) as a sarcolemma marker and DAPI (blue) for nuclei visualisation. A magnified view of the sections shows the centrally nucleated fibers (CNF) proportion in two regions of the section. Tiled images were taken at 10× magnification and stitched together using the LAS X software. Scale bars represent 100 µm. (**B**) CNF as a proportion of total myofibers analysed per TA section from *mdx52-Xist*^Δhs^ animals expressing low, medium, and high dystrophin levels (*n*=3). A single *mdx52* animal was included as a reference. (**C**) Correlation analysis of dystrophin percentage and average CNF percentage in *mdx52-Xist*^Δhs^ TA sections, with a single *mdx52* animal as a reference. Dystrophin protein percentage was derived from western blot analysis. (**D**) Myofiber size variability in TA sections of *mdx52-Xist*^Δhs^ animals (*n*=3) visualized through proportion of fibers of specific cross-sectional area (CSA). Values are mean±SD. Statistical significance was assessed by one-way ANOVA with Bonferroni *post hoc* test relative to the Low dystrophin expressing group, \*\**P*<0.01. Nuclei were stained with DAPI.

### Dystrophin patchiness is maintained in aged *mdx52-Xist*^Δhs^ muscle

It has been proposed that dystrophin protein may accumulate with time following CRISPR-Cas9-mediated correction as a result of positive selection of corrected, dystrophin-expressing myofibers.^4,5,10,21^ To investigate this dystrophin accumulation phenomenon, we generated a separate cohort of *mdx52-Xist*^Δhs^ female F1 mice (*N*=21) and sacrificed them at 60 weeks of age. Dystrophin expression was determined in TA muscles by western blot in this ‘aged’ cohort and animals retrospectively assigned to high (53-91% of wild-type dystrophin, *n*=5), medium (21-46%, *n*=8), and low (3-20%, *n*=8) dystrophin expression as described above (**Figure 4A-C**). The mean dystrophin expression value for all aged *mdx52-Xist*^Δhs^ animals was ~2.8-fold higher than the mean of all 6-week-old *mdx52-Xist*^Δhs^ animals (*P*<0.001), consistent with the enrichment of dystrophin positive myofibers as a consequence of positive selection. However, the patchy pattern of sarcolemmal dystrophin expression was maintained in aged animals at all dystrophin expression levels (**Figure 4C**). Analysis of isolated single EDL myofibers from aged *mdx52-Xist*^Δhs^ showed that dystrophin, β-dystroglycan (DAG1), and α-dystrobrevin (DTNA) exhibited ‘zebra-like’ patchy immunostaining patterns, while this effect was much less clear for nNOS (**Figure 5**), similar to those observations in adult animals (**Figure 2**).

**Figure 4.**
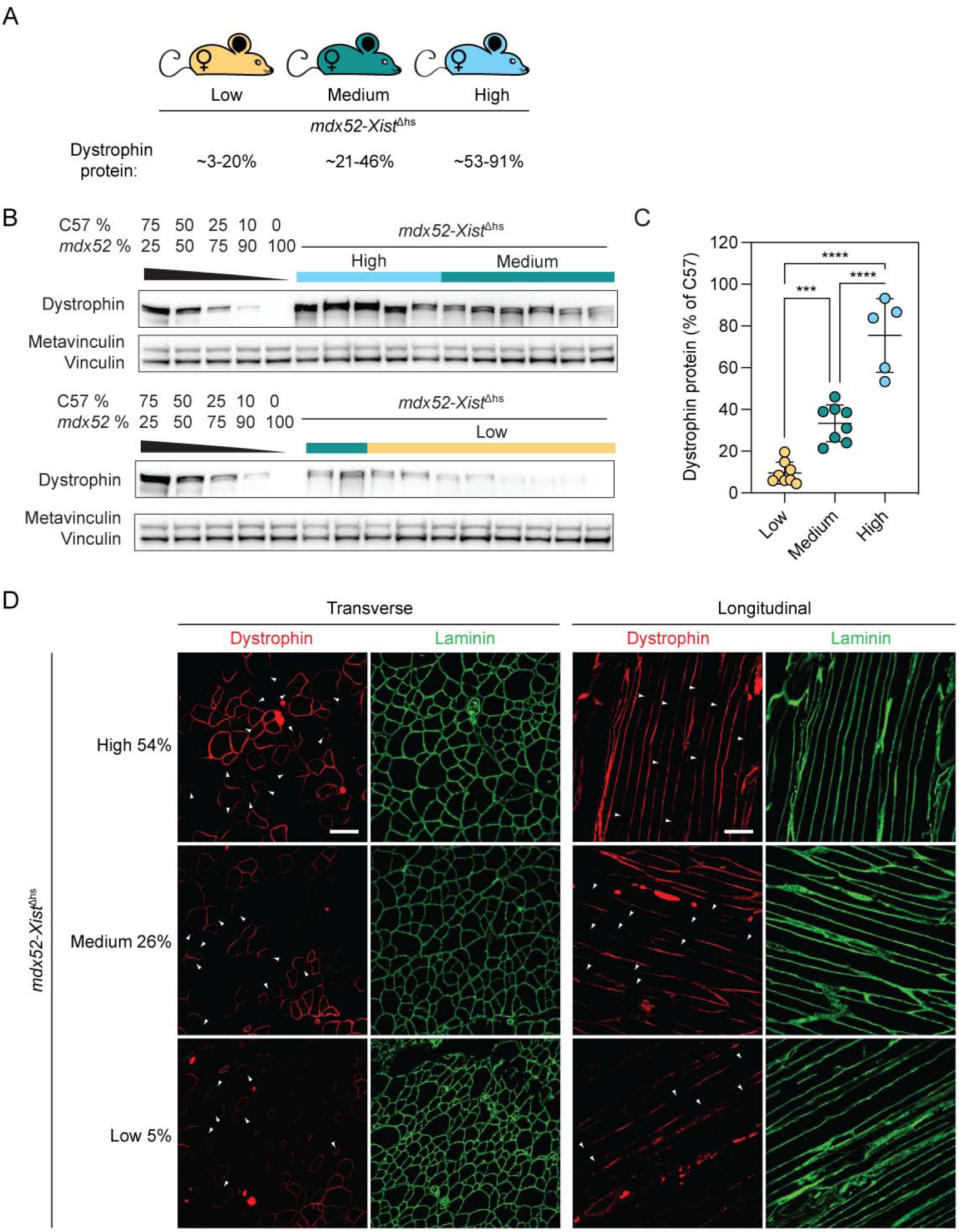
Dystrophin patchiness is maintained in aged *mdx52-Xist*^Δhs^ muscle. (**A**) Aged *mdx52-Xist*^Δhs^ animals were assigned to high (*n*=5), medium (*n*=8), and low (*n*=8) dystrophin expressing groups post-mortem based on protein quantification in 60-week-old TA muscles as determined by (**B**) Western blot analysis. (**C**) Dystrophin was quantified by comparison to standard curves containing defined mixtures of C57 and *mdx52* TA lysates. Vinculin was utilised as a loading control. (**D**) Representative immunofluorescence staining of dystrophin and laminin in transverse and longitudinal TA muscle sections of 60-week-old *mdx52-Xist*^Δhs^ animals from high, medium, and low dystrophin-expressing groups. Within-fiber, patchy dystrophin expression resulting from skewed X-chromosome inactivation indicated with arrowheads. Scale bars indicate 100 µm, images taken at 20× magnification. The percentage values indicate total dystrophin quantification in the animals from which the sections were derived. Values are mean±SD. Statistical significance was assessed by one-way ANOVA with Bonferroni *post hoc* test, \*\*\**P*<0.001, \*\*\*\**P*<0.0001.

**Figure 5.**
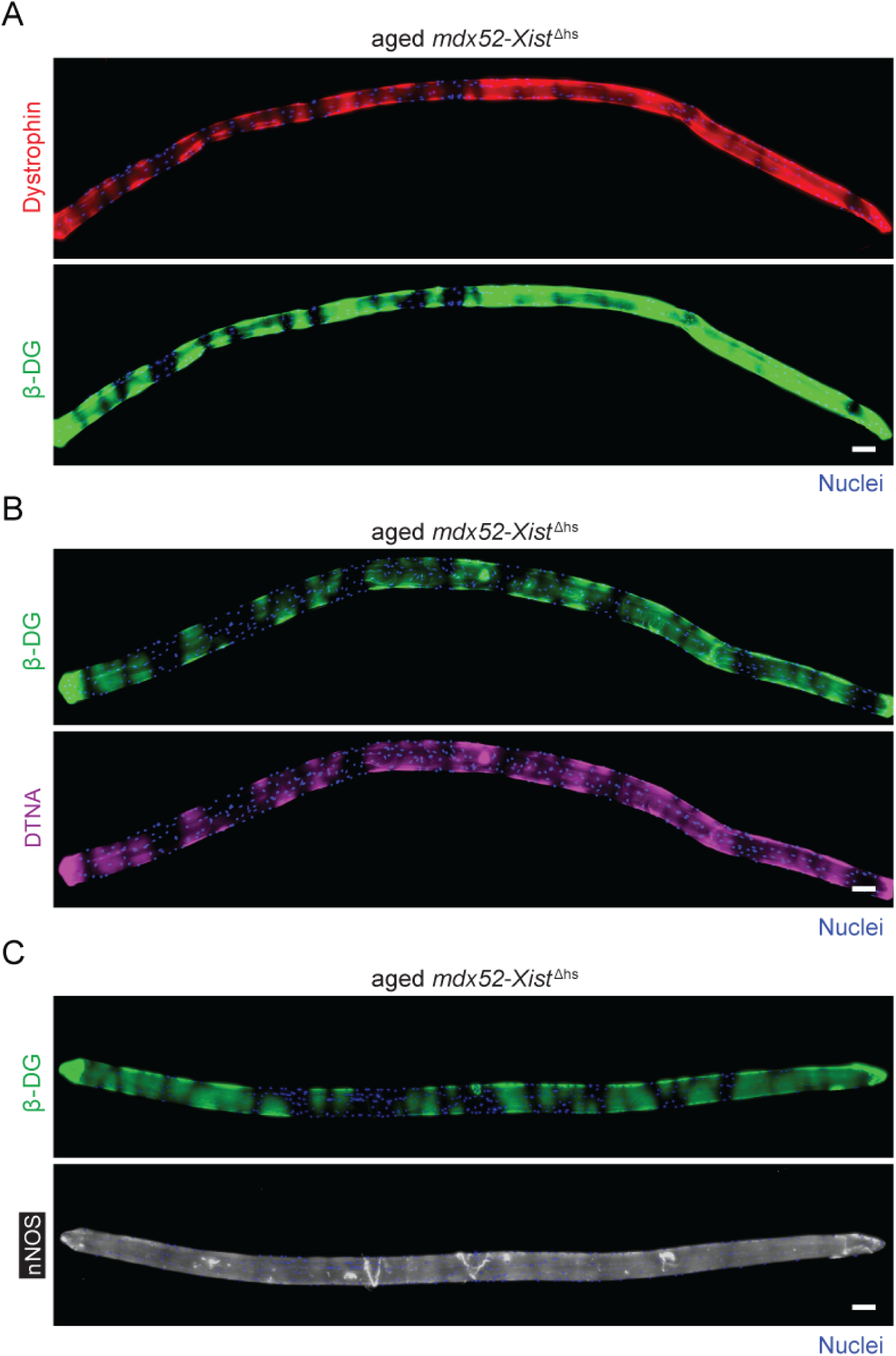
Dystrophin patchiness is maintained in aged *mdx52-Xist*^Δhs^ isolated myofibers. Representative immunofluorescence co-staining of (**A**) dystrophin and β-DG, (**B**) β-DG and DTNA, and (**C**) β-DG and nNOS in isolated single 60-week-old *mdx52-Xist*^Δhs^ EDL myofibers. Tiled images were taken at 10× magnification and stitched together in LAS X software. Scale bars indicate 100 µm. Nuclei were stained with DAPI.

### Dystrophin expression is inversely correlated with muscle histopathology in aged *mdx52-Xist*^Δhs^ muscles

Histopathological analysis in aged *mdx52-Xist*^Δhs^ TA muscles revealed abundant CNFs and foci of small diameter regenerating fibers (**Figure 6A**) at all levels of dystrophin expression. Mean CNF values were much larger than in the 6-weeks-old *mdx52-Xist*^Δhs^ mice with 35.9%, 51.1%, and 57.1% for high, medium, and low dystrophin expressing muscles, respectively (**Figure 6B**). The percentage of CNFs was inversely correlated with dystrophin expression (Spearman’s *r*=-0.5099, *P*=0.0257, **Figure 6C**), although the correlation was substantially weaker than that observed for 6-week-old animals (**Figure 3C**). Analysis of myofiber cross-sectional area revealed no differences between *mdx52-Xist*^Δhs^ groups (**Figure 6D**). Together, these results suggest that muscles expressing dystrophin in a patchy manner continue to degenerate and regenerate throughout life, and that these pathological processes are to some extent ameliorated with higher levels of dystrophin expression.

**Figure 6.**
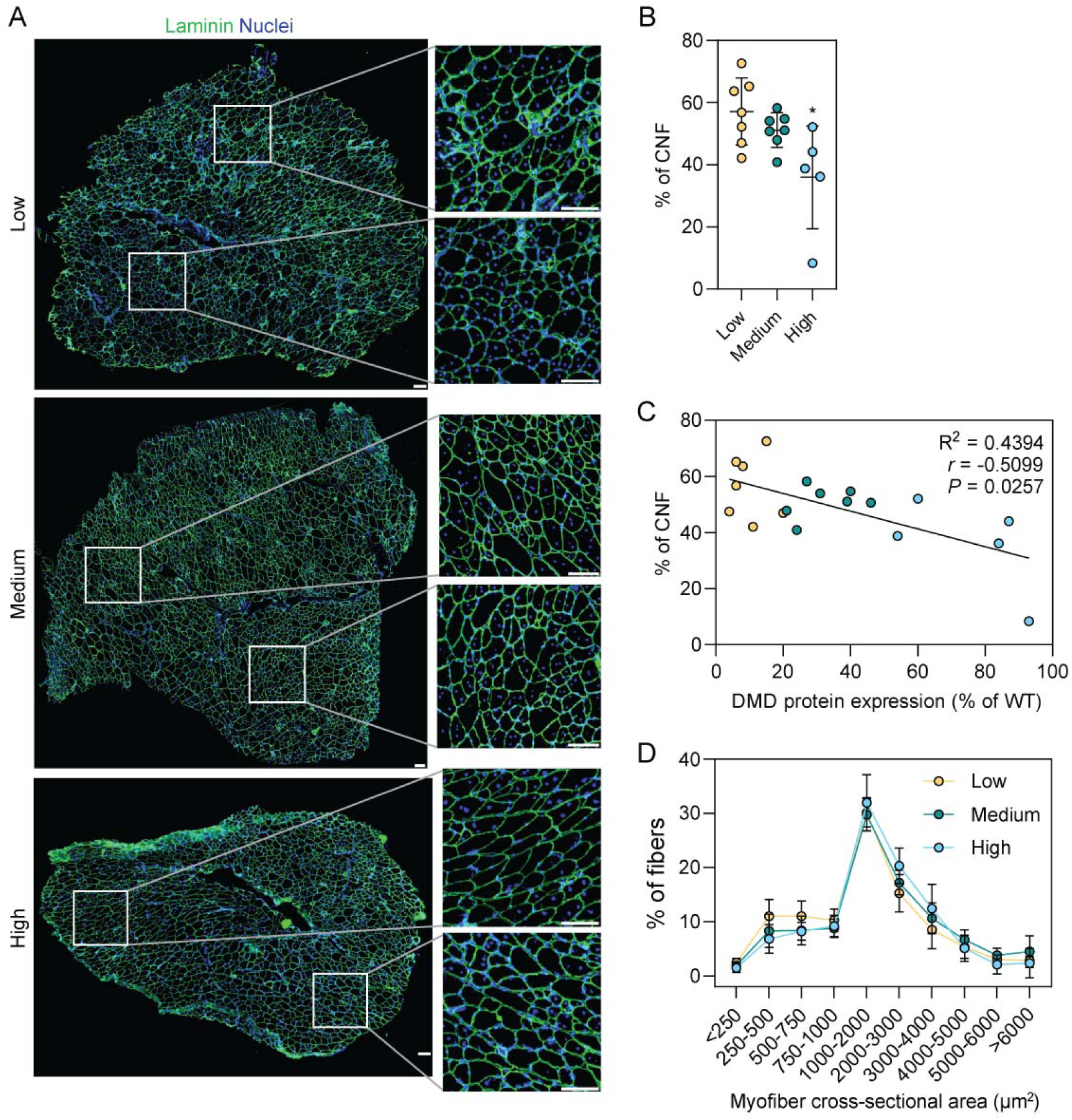
Dystrophin expression is inversely correlated with histopathology in aged *mdx52-Xist*^Δhs^ muscle sections. (**A**) Representative immunofluorescence images of transverse TA muscle sections stained from aged (60-week-old) *mdx52-Xist*^Δhs^ animals expressing high (*n*=5), medium (*n*=7), and low (*n*=7) dystrophin levels. Sections were stained for laminin (green) as a sarcolemma marker and DAPI (blue) for nuclei visualisation. A magnified view of the sections shows the centrally nucleated fibers (CNF) proportion in two regions of the section. Tiled images were taken at 10× magnification and stitched together using the LAS X software. Scale bars represent 100 µm. (**B**) CNF as a proportion of total myofibers analysed per TA section. (**C**) Correlation analysis of dystrophin percentage and average CNF percentage in analysed *mdx52-Xist*^Δhs^ TA sections. (**D**) Myofiber size variability in TA sections of *mdx52-Xist*^Δhs^ animals visualised through proportion of fibers of specific cross-sectional area (CNS). Values are mean±SD. Statistical significance was assessed by one-way ANOVA with Bonferroni *post hoc* test relative to the Low dystrophin expressing group, \**P*<0.05. Nuclei were stained with DAPI.

### Utrophin expression does not correlate with patchy dystrophin levels in adult and aged *mdx52-Xist*^Δhs^ animals

Utrophin and dystrophin exhibit reciprocal expression patterns during muscle development and dystrophic pathology.^22^ Moreover, when expressed together, they bind to the same sites at the sarcolemma, as evidenced by electron microscopy detection of both proteins in muscles of transgenic mice.^23^ Due to the dystrophic nature of *mdx52-Xist*^Δhs^ myofibers, and the presence of dystrophin in positive myonuclear domains, unrestricted availability of binding sites is expected within dystrophin-negative regions. We therefore reasoned that utrophin and dystrophin might exhibit reciprocal patterns of myonuclear domain restriction.

To test this hypothesis, utrophin western blot was performed on muscle lysates from adult and aged *mdx52-Xist*^Δhs^ animals (**Figure S2A,B**). Utrophin expression was variable but detected in all analysed animals, regardless of age or dystrophin levels. Expression of utrophin and dystrophin in TA muscles was not correlated for either 6-week-old or aged animals (Spearman’s *r* = −0.2807, *P* = 0.6604, and *r* = −0.03506, *P* = 0.3023, respectively, **Figure S2C,D**).

To assess the localization of utrophin in patchy dystrophin muscles, utrophin immunofluorescence was performed in transverse TA sections from *mdx52-Xist*^Δhs^ animals identified as exhibiting high utrophin expression within the low dystrophin expressing group (**Figure S2E**). 12-week-old *mdx52* tissue sections were used as control whereby a clear utrophin signal was detected at the membrane of relatively small, centrally-nucleated myofibers organized in densely packed clusters (**Figure S2E**). Accordingly, several small groups of newly formed CNFs with positive utrophin staining were detected in aged *mdx52-Xist*^Δhs^ muscle expressing low levels of dystrophin. Notably, no utrophin expression was observed at the sarcolemma of larger *mdx52-Xist*^Δhs^ TA myofibers (**Figure S2E**). As such, utrophin expression in *mdx52-Xist*^Δhs^ animals reflects the degree of ongoing muscle regeneration and is not concentrated in dystrophin-negative myonuclear domains.

Serum microRNAs (miRNAs) have been investigated as minimally-invasive biomarkers in the context of DMD.^24^ In particular, we have previously reported that serum myomiR levels are inversely correlated with dystrophin expression levels following antisense oligonucleotide-mediated exon skipping, suggesting that they may constitute promising pharmacodynamic biomarkers.^8,25^ In 6-week-old animals, myomiRs were inversely correlated with dystrophin expression level (**Figure S3A-F**). Conversely, in aged animals, there was no difference between serum myomiR levels between *mdx52-Xist*^Δhs^ animals, and accordingly no correlation with dystrophin protein expression (**Figure S3G-L**). These observations are consistent with our previous findings that serum myomiR levels are associated with regenerative pathology, which declines with age, and is effectively absent in aged animals.^26,27^

### Dystrophin is not expressed in centrally-nucleated *mdx52-Xist*^Δhs^ myofiber segments

Inspection of single isolated myofibers revealed the existence of three types of fiber based on the degree of central nucleation; (i) non-centrally-nucleated (59.9%), (ii) uniformly centrally-nucleated (16.6%), and (iii) segmented centrally-nucleated, whereby chains of centrally-located myonuclei were restricted to regions within the associated myofiber (23.5%) (**Figure 7A**). Non-centrally nucleated myofibers overwhelmingly (99.6%) exhibited patchy, ‘zebra-like’ patterns of dystrophin distribution (**Figure 7B**, and similar to micrographs in **Figures 2 and 5A**). We next classified segmented fibers according to dystrophin/β-dystroglycan (i.e. DAPC) expression, with signal in the non-centrally-nucleated region only (75.4%), coverage in both segments (18.4%), or no dystrophin/DAPC expression present at all (6.2%) (**Figure 7C**). Similarly, centrally-nucleated myofibers and myofiber segments were found to be almost completely devoid of dystrophin/β-dystroglycan expression. In fully centrally-nucleated myofibers, 88% contained no dystrophin or β-dystroglycan (i.e. DAPC) expression. The remaining 12% of myofibers exhibited some expression, although this was frequently limited to very small regions of membrane (**Figure 7D**). Importantly, in DAPC-positive fully-centrally-nucleated fibres, none exhibited the patchy, ‘zebra-like’ staining pattern. This finding suggests that the observed dystrophin absence in centrally-nucleated regions is very unlikely to be driven by an XCI effect associated with the *Xist*^Δhs^ model.

**Figure 7.**
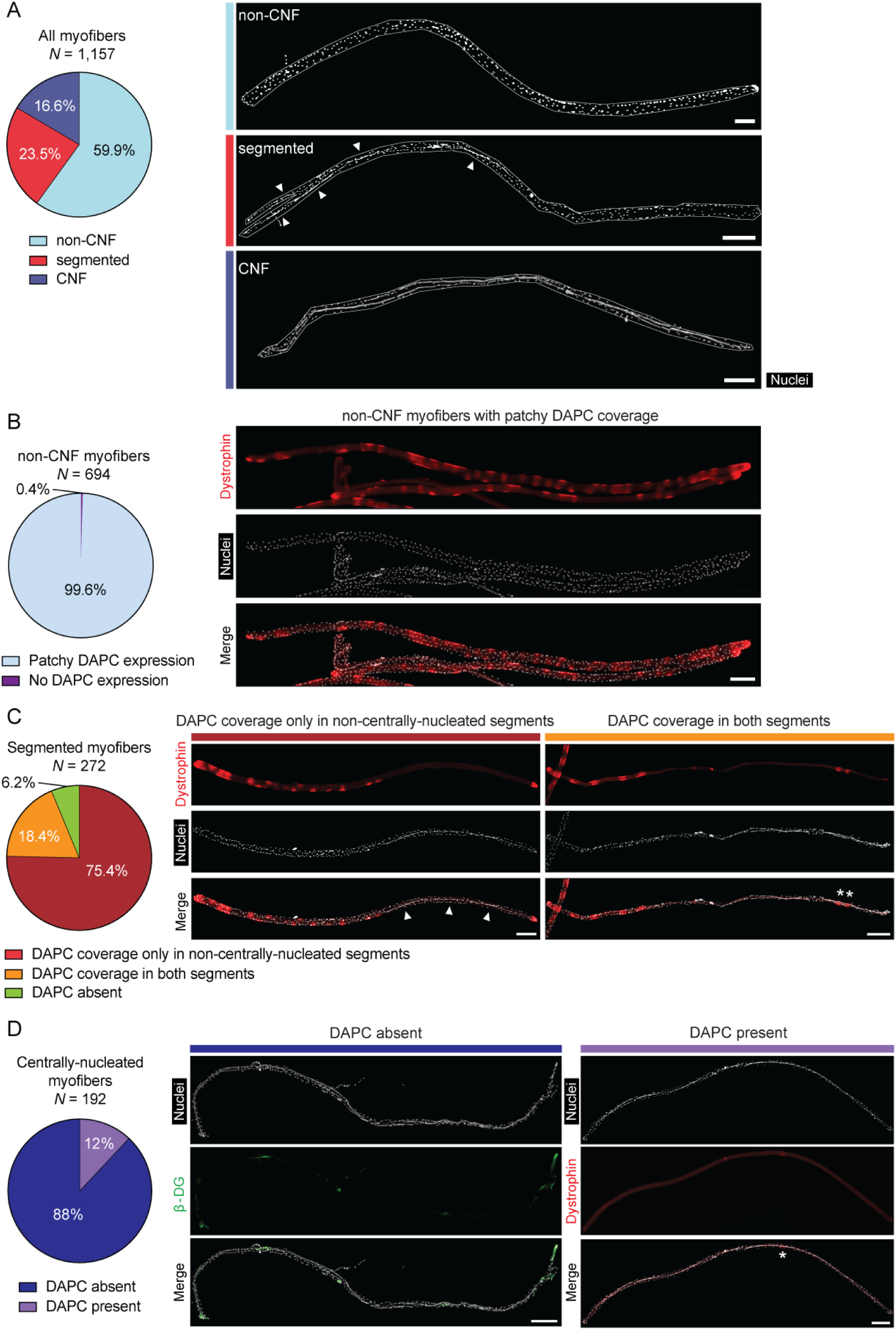
Dystrophin is largely absent in centrally-nucleated myofibers/fiber segments. (**A**) Single isolated EDL myofibers harvested from adult (12-17 week old) *mdx52-Xist*^Δhs^ mice (*N*=1,157) were sorted into non-centrally-nucleated (non-CNF), segmented (i.e. containing both centrally-nucleated and non-centrally-nucleated fiber regions), and fully centrally-nucleated (CNF) categories. (**B**) Non-CNFs (*N*=694) were further classified into those with a ‘zebra-like’ patchy DAPC pattern of expression, and those with no DAPC expression. (**C**) Segmented myofibers (*N*=272) were further classified based on whether DAPC proteins (i.e. dystrophin or β-dystroglycan) were detected in non-CNF segments, both CNF and non-CNF segments, or absent from both segments. (**D**) Fully centrally-nucleated myofibers (*N*=192) were classified based on whether or not DAPC proteins were detected. Arrow heads indicate centrally-nucleated regions-of-interest. Asterisks indicate DAPC positive regions-of-interest. Tiled images were acquired at 10× magnification and stitched using the LAS X software. Scale bars indicate 200 µm. Nuclei were stained with DAPI.

The absence of dystrophin and β-dystroglycan expression in centrally-nucleated myofiber segments was even more apparent in segmented myofibers from aged (60-week-old) *mdx52-Xist*^Δhs^ animals (**Figure 8A**), with aged fully-CNF myofibers being largely devoid of DAPC protein expression (**Figure 8B**). A representative bulk preparation of myofibers from a single 60-week-old *mdx52*-*Xist*^Δhs^ illustrating this point is shown in **Figure S4**. Notably, the aged myofibers were evidently hypertrophic and frequently contained multiple chains of centrally-located myonuclei. These observations indicate that centrally-nucleated myofibers in *mdx52-Xist*^Δhs^ mice at this age are not recently regenerated. Immunostaining for the nuclear envelope marker Lamin B1 (LAMB1) showed that these nuclei chains consist of intact myonuclei squashed together, rather than fused together (**Figure S5**).

**Figure 8.**
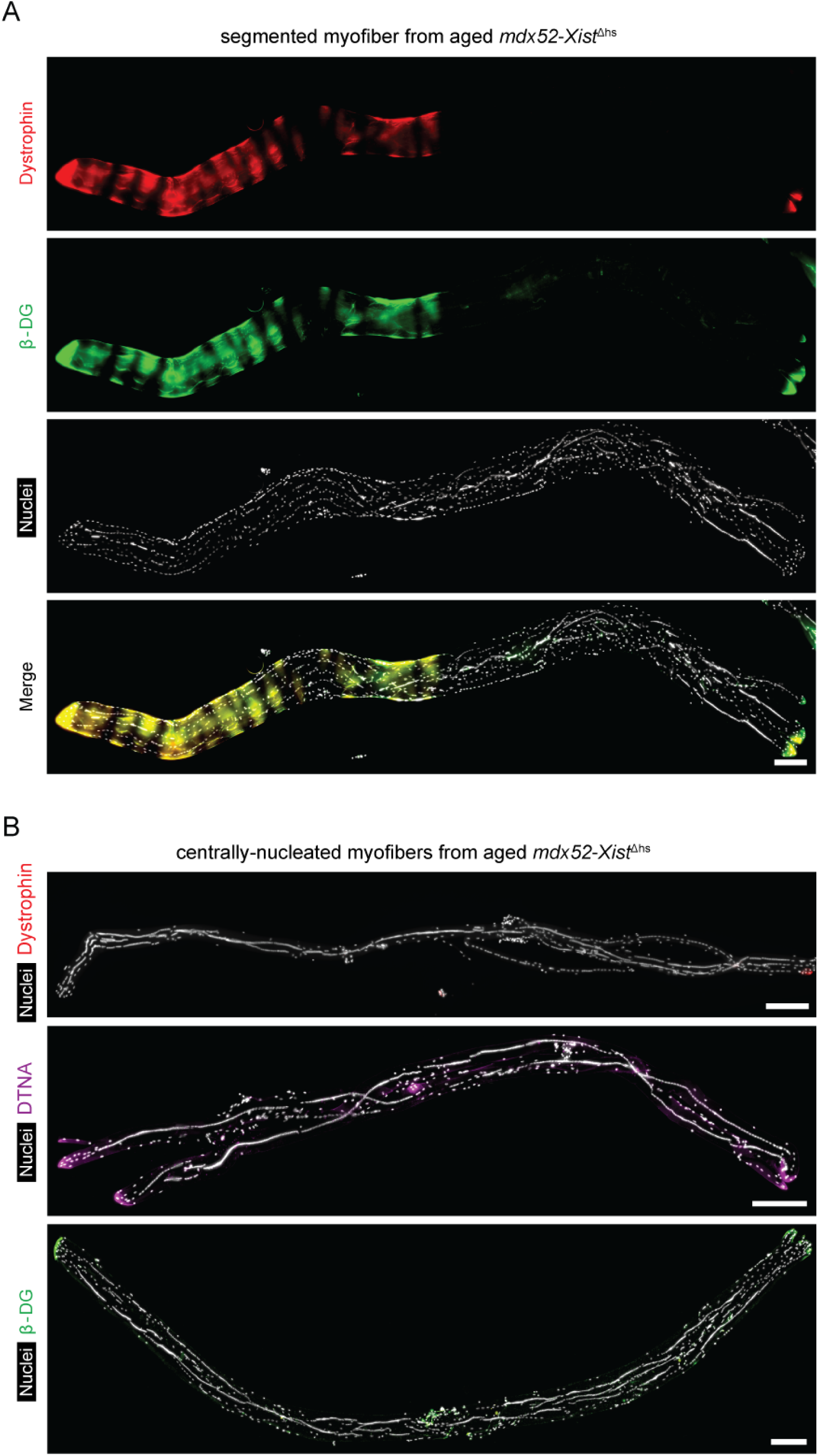
Dystrophin and DAPC protein expression is largely absent in centrally-nucleated myofibers/fiber segments in aged *mdx52-Xist*^Δhs^ mice. Single EDL myofibers isolated from 60-week-old *mdx52-Xist*^Δhs^ mice were analysed by immunofluorescence and representative micrographs shown for (**A**) dystrophin and β-dystroglycan (β-DG) co-staining in segmented myofibers (i.e. containing both centrally-nucleated and non-centrally-nucleated regions), and (**B**) dystrophin, α-dystrobrevin (DTNA) and β-DG in fully centrally-nucleated myofibers. Tiled images were acquired at 10× magnification and stitched using the LAS X software. Scale bars represent 200 µm. Nuclei were stained with DAPI.

To assess whether muscle injury alone could induce a similar impairment in dystrophin expression, we injected adult wild-type mice (20-30-week-old, both males and females) with the muscle toxicant BaCl_2_ in order to induce acute myonecrosis and regeneration. Animals were harvested after 29 days, after which the muscle morphology was restored but central nucleation persisted. Immunofluorescence staining in these animals revealed complete sarcolemmal dystrophin coverage, including in centrally-nucleated myofibers, suggesting that muscle regeneration *per se* does not recapitulate the phenomenon of dystrophin absence in centrally-nucleated myofiber segments observed in *mdx52-Xist*^Δhs^ mice (**Figure S6**).

Taken together, these data show that dystrophin/the DAPC is largely absent in CNF myofibers and in the centrally-nucleated regions of segmented myofibers. The absence of a similar effect in injured wild-type muscle suggests that this phenomenon is a feature of dystrophic muscle.

### Translation of dystrophin is specifically impaired in centrally-nucleated myofiber segments

The absence of dystrophin in centrally-nucleated myofiber segments might be the result of a global impairment in either transcription or translation. RNA-FISH analysis using a pool of probes spanning the *Dmd* transcript revealed puncta evenly distributed throughout *mdx52-Xist*^Δhs^ single isolated segmented myofibers independent of dystrophin expression (**Figure 9A**). (Notably, this assay is not capable of distinguishing between the mutant and wild-type *Dmd* alleles). Analysis of centrally-nucleated *mdx52-Xist*^Δhs^ single isolated myofibers showed that both *Dmd* transcripts and titin (TTN) protein were uniformly distributed throughout all myofibers assessed (**Figure 9B**). Moreover, TTN exhibited a characteristic pattern of sarcomeric striation, indicative of myofiber maturity. TTN and filamentous Actin (F-Actin) were found to be evenly distributed throughout both centrally-nucleated and non-centrally-nucleated myofiber regions, in stark contrast to the pattern observed for dystrophin (**Figure 9C**). These data demonstrate that there is no shortage or mislocalization of *Dmd* mRNAs in centrally-nucleated myofiber regions, and that there is no local impairment in global protein translation. As such, these data suggest that dystrophin protein expression is specifically impaired in *mdx52-Xist*^Δhs^ centrally-nucleated myofiber regions at the level of translation.

**Figure 9.**
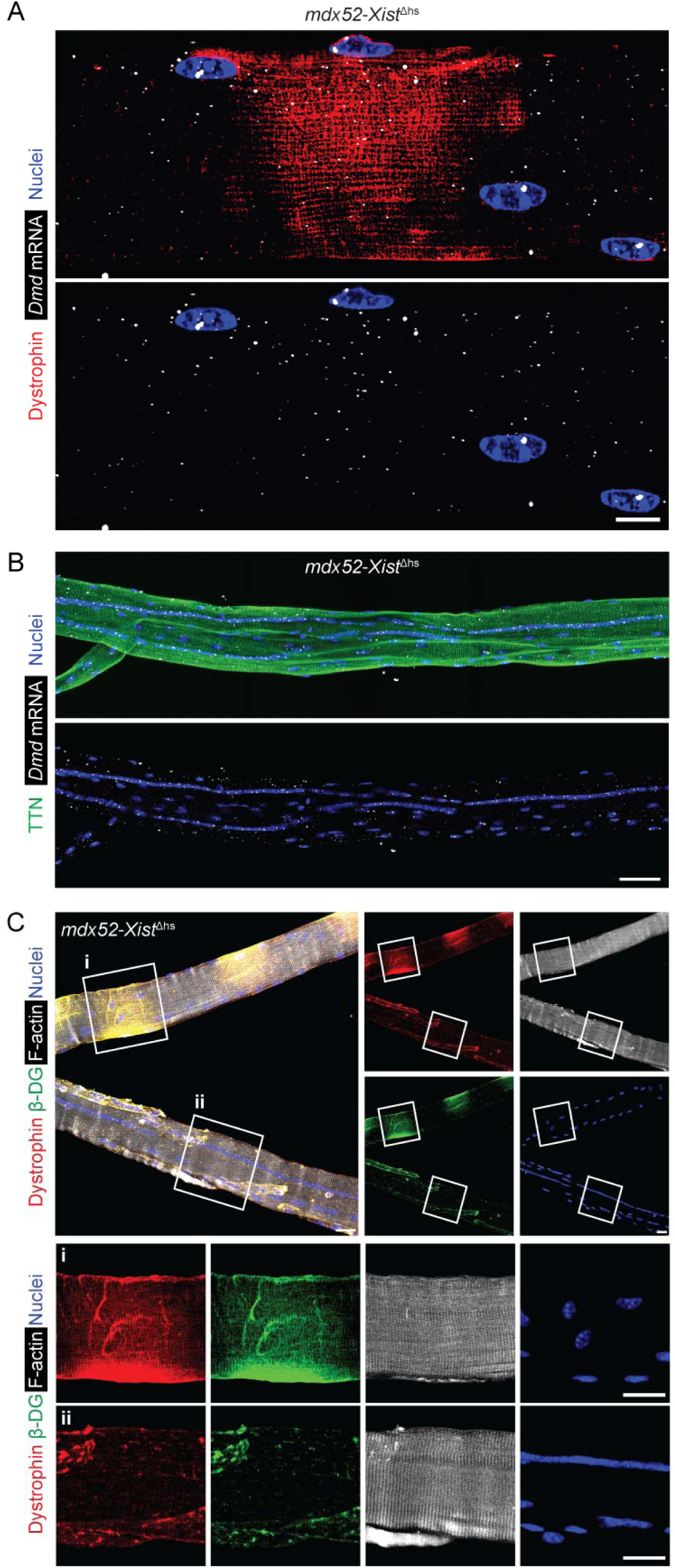
Dystrophin translation is suppressed in *mdx52-Xist*^Δhs^ centrally-nucleated myofiber segments. Single EDL myofibers were isolated from adult (8-week-old) *mdx52-Xist*^Δhs^ and analysed for immunofluorescence and RNA fluorescence *in situ* hybridization (FISH). (**A**) Representative micrograph of a single isolated EDL myofiber from an *mdx52-Xist*^Δhs^ animal (12-week-old) showing combined immunostaining for dystrophin protein and HCR-FISH for *Dmd* mRNA. Image taken at 40× magnification, scale bar represents 10 µm. (**B**) Representative micrograph of a centrally-nucleated myofiber stained for TTN protein and dystrophin mRNA. Tiled images were acquired at 25× magnification and stitched together using the ZEN Blue software. Scale bar represents 50 µm. (**C**) Representative micrograph of a segmented myofiber (i.e. containing both centrally-nucleated and non-centrally-nucleated regions) and a fully centrally-nucleated myofiber in the same frame stained for dystrophin, β-dystroglycan (β-DG), and filamentous-actin (F-actin). Selected regions showing (**i**) a patchy, non-centrally nucleated segment, and (**ii**) a centrally-nucleated segment, are enlarged and shown inset. Images were acquired at 25× magnification. Scale bars represent 20 µm. Nuclei were stained with DAPI.

### Microtubule network disruption is similar in dystrophin positive and negative segments

The absence of dystrophin expression in centrally-nucleated regions might be a consequence of impaired mRNA trafficking following microtubule network disruption. One of the hallmarks of correct myofiber organization is the intricately organized microtubule network, which was recently shown to facilitate the active transport of various RNAs and proteins, including the ribosomal machinery, throughout the cell.^28,29^ Notably, dystrophin protein contains a microtubule-binding domain and therefore has been proposed to stabilize the myofiber microtubule cytoskeleton.^30,31^ This role is supported by the fact that the microtubule network is significantly disorganised in dystrophic mice, with costameric (transverse) components being the most severely affected.^30,32^ Thus, it is likely that non-uniformly distributed dystrophin can modify the organization of the microtubules in *mdx52-Xist*^Δhs^ mice, thereby partially facilitating correct mRNA transcript trafficking. TeDT (texture detection technique) analysis of microtubules was performed on images acquired from *mdx52-Xist*^Δhs^, *mdx52*, and wild-type C57 EDL myofibers (*n*=40, 31, and 32 regions of interest, respectively). The characteristic peak at the 90° intersection angle (representing the transverse microtubules) together with a high vertical directionality score was detected in adult wild-type C57 myofibers (**Figure 10A-C**). In agreement with previous reports, the microtubule network was visibly disorganised in *mdx52* animals, with a corresponding significant loss of transverse microtubules (**Figure 10B,C**).^30,32^ This disorganised pattern was partially restored in *mdx52-Xist*^Δhs^ myofibers as represented by an intermediate distribution of microtubule intersection angles and vertical directionality scores (**Figure 10B,C**). These results show that non-uniformly distributed dystrophin in the *mdx52-Xist*^Δhs^ model is associated with an intermediately distorted microtubule network.

**Figure 10.**
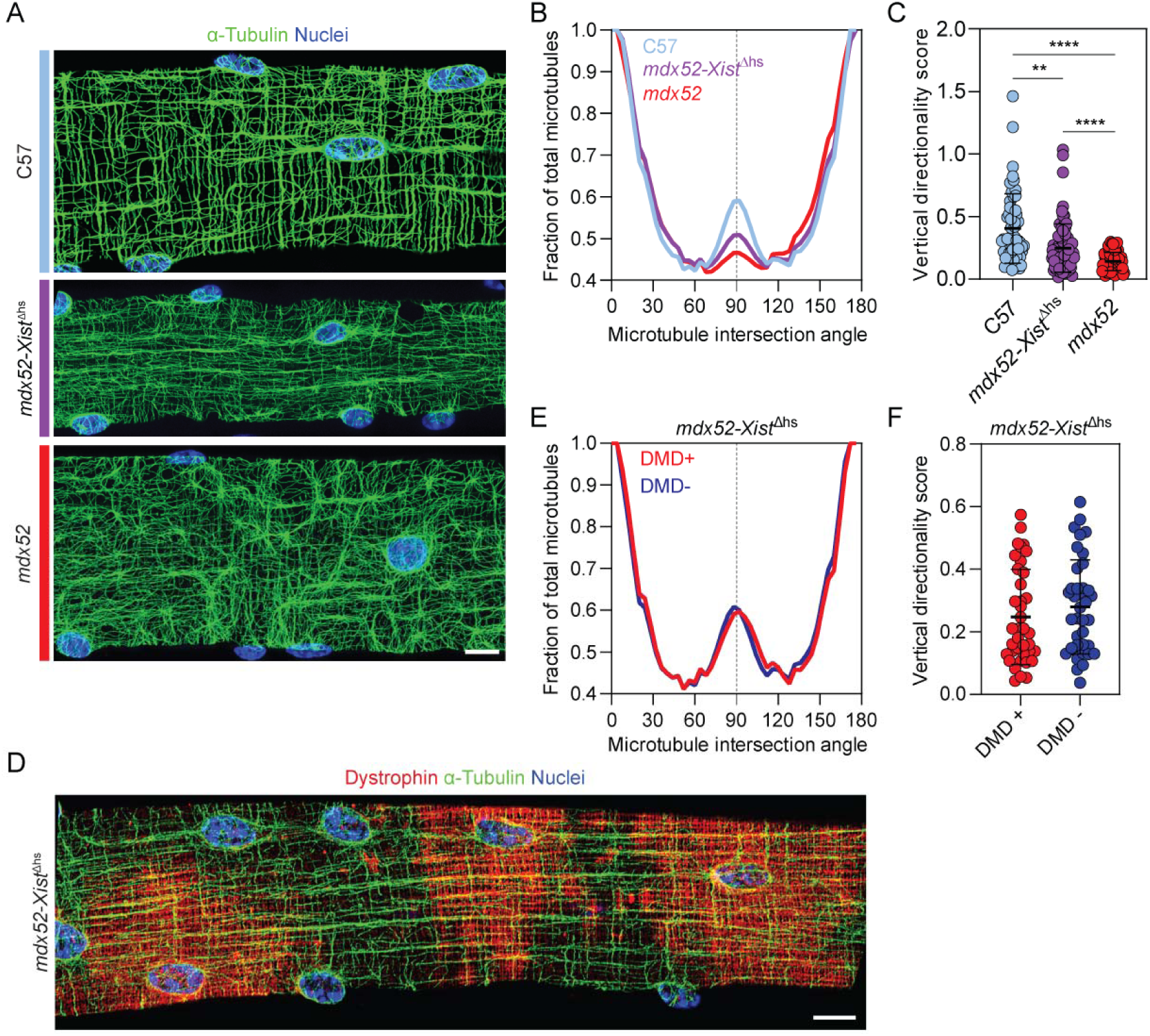
*mdx52-Xist*^Δhs^ myofibers exhibit intermediate microtubule network disorganization that is similar in dystrophin positive and negative segments. (**A**) Representative micrographs of immunostaining for α-tubulin to show cortical microtubule network organization in adult (8-12-weeks-old) C57 wild type, *mdx52*, and *mdx52-Xist*^Δhs^ single isolated EDL myofibers. (**B**) Histogram of mean distribution of microtubules of different intersection angles relative to myofiber long axis. The transverse, costameric microtubule peak (90°) is marked with a dotted line. (**C**) Vertical directionality scores reflecting the summed values of microtubules present between 80° to 100° within each myofiber. (Sample sizes are; C57: *n*=60 ROIs, derived from 32 myofibers, *mdx52*: *n*=64 ROIs, derived from 31 myofibers, *mdx52-Xist*^Δhs^: *n*=79 ROIs, derived from 40 myofibers). (**D**) Representative micrographs of co-immunostaining for α-tubulin and dystrophin to show cortical microtubule network organization in a patchy dystrophin *mdx52-Xist*^Δhs^ single isolated EDL myofiber. (**E**) Histogram of mean distribution of microtubules of different intersection angles relative to myofiber long axis in dystrophin positive, and negative myonuclear domains of *mdx52-Xist*^Δhs^ EDL myofibers. The transverse, costameric microtubule peak (90°) is marked with a dotted line. (Sample sizes are; DMD+: *n*=42 ROIs, derived from 24 myofibers, DMD-: *n*=40 ROIs, derived from 25 myofibers). (**F**) Vertical directionality score reflecting the summed values of microtubules present between 80 to 100 degrees within each fiber. Images taken at 40× magnification, scale bars represent 10 µm. Plotted values are mean±SD. Statistically significant differences were assessed by one-way ANOVA with Bonferroni *post hoc* test or two-tailed Student’s *t*-test, as appropriate. \*\**P*<0.01, **** *P*<0.0001. Nuclei were stained with DAPI.

We were next motivated to determine whether there was a difference between microtubule network organization in dystrophin-positive and -negative *mdx52-Xist*^Δhs^ myofiber segments (**Figure 10D-F**). No difference in microtubule lattice organization was observed between the analysed domains in terms of microtubule intersection angle distribution (**Figure 10E**) or vertical directionality scores (**Figure 10F**). This suggests that the local absence of dystrophin alone may not be sufficient to induce cytoskeletal network disruption.

### Aged *mdx52-Xist*^Δhs^ animals contain high proportions of hypertrophic centrally-nucleated myofibers in the absence of active regeneration

Central nucleation is associated with muscle regeneration, but is known to persist long after injury in mice (as long as 21 months).^33,34^ In addition, regenerating myofibers exhibit small cross-sectional areas and are positive for development-associated markers such as embryonic myosin heavy chain and utrophin.^35^ To better understand the phenomenon of dystrophin absence in centrally nucleated myofibers/fiber segments, we analysed the extent of central nucleation in adult (6 week) and aged (60 week) *mdx52-Xist*^Δhs^ TA muscle sections.

Central nucleation was shown to significantly increase in TA muscle sections from aged animals (**Figure 11A**). Furthermore, there was a shift towards a greater number of central myonuclear chains with age in isolated EDL myofibers (**Figure 11B**). Analysis of myofiber cross-sectional area revealed that aged *mdx52-Xist*^Δhs^ animals exhibited a pronounced shift towards larger fibers (**Figure 11C**), and that there was a statistically significant (*P*<0.0001) shift in the mean Feret diameter in centrally-nucleated fibers (**Figure 11D**). Taken together, these data show that CNFs undergo substantial hypertrophy with age, which is likely driven by the progressive accretion of new myonuclei. Indeed, the number of nuclei per myofiber volume was increased in CNF regions compared with non-CNF regions in *mdx52* mice (**Figure S7**).^36,37^ These observations, together with the limited dispersion of central nuclei to the myofiber periphery observed in mice,^33,34^ demonstrate that CNFs at later stages largely do not constitute a population of recently formed muscle cells.^34,38^ Together, these results emphasise the fact that the proportion of CNF at later stages of life in mice reflects the cumulative history of regeneration, and not recently regenerating immature muscle.^38^ The latter point is important, because myofiber immaturity could be a potential explanation for the absence of dystrophin in centrally-nucleated myofiber regions. Furthermore, the expression of late-stage markers of muscle maturity (i.e. TTN, **Figure 9B**) and the absence markers of early-stage muscle development (i.e. UTRN, **Figure S2**) in CNFs lends further credence to the notion that these myofibers are not recently regenerating and immature.

**Figure 11.**
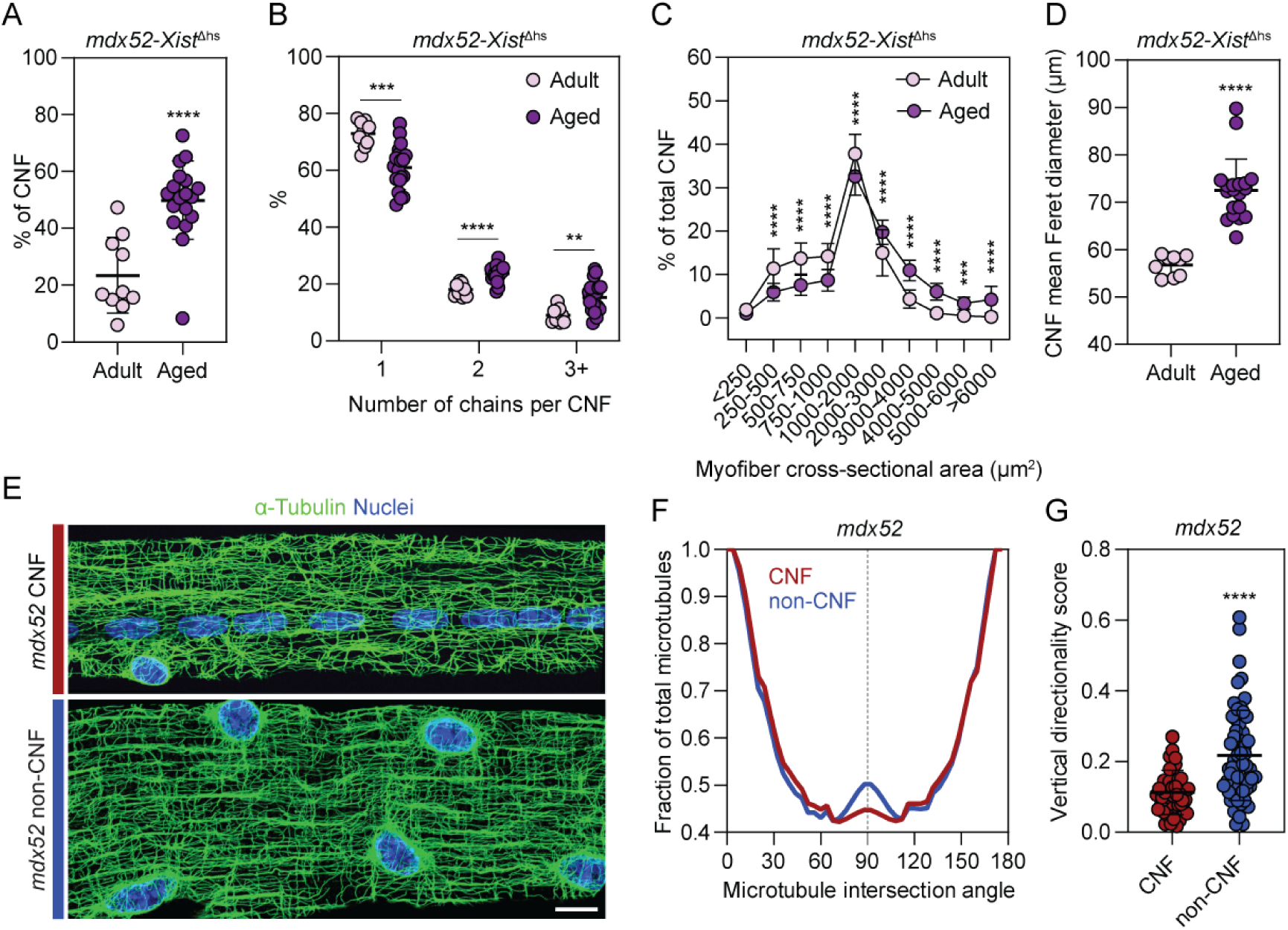
Central nucleation accumulates with age in *mdx52-Xist*^Δhs^ mice and is associated with microtubule network disruption. Adult (6-week-old) and aged (60-week-old) *mdx52-Xist*^Δhs^ TA muscle sections were compared for (**A**) the percentage of CNF myofibers, (**B**) the number of chains per centrally-nucleated myofiber, (**C**) mean Feret diameter of CNFs, and (**D**) Distribution of CNF cross-sectional area (CSA). Separately, adult (12-week-old) *mdx52* single isolated EDL myofibers were harvested and analysed for microtubule network organization in centrally-nucleated and non-centrally-nucleated myofibers. (**E**) Representative micrographs of immunostaining for α-tubulin. (**F**) Histogram of mean distribution of microtubules of different intersection angles relative to myofiber long axis in centrally-nucleated and non-centrally-nucleated *mdx52* myofibers. The transverse, costameric microtubule peak (90°) is marked with a dotted line. (**G**) Vertical directionality score reflecting the summed values of microtubules present between 80 to 100 degrees within each fiber. (Sample sizes are; CNF: *n*=44 ROIs, derived from 22 myofibers, non-CNF: *n*=60 ROIs, derived from 27 myofibers). Images taken at 40× magnification, scale bar represents 10 µm. Values are mean±SD. Statistically significant differences were assessed by Student’s *t*-test, \*\**P*<0.01, \*\*\**P*<0.001, \*\*\*\**P*<0.0001. Nuclei were stained with DAPI.

Notably, in mature myofibers, nuclei play a crucial role as microtubule organization centers. ^39,40^ As such, the distinct localization of myonuclei within CNFs and non-CNFs could potentially affect the organization of the microtubule network. To assess the contribution of central-nucleation to microtubule organization, cortical microtubules were analysed in CNF and non-CNF EDL myofibers harvested from 12-week-old *mdx52* mice, as these muscles are expected to contain very high proportions of centrally-nucleated myofibers.^38^ The microtubule network was visibly disorganized in CNFs (**Figure 11E**), which was accompanied by a quantitative decrease in transverse microtubules (**Figure 11F**) and vertical directionality scores in CNFs (*P*<0.0001) (**Figure 11G**). These results show, that within dystrophic myofibers, the microtubule lattice is substantially more disrupted in centrally-nucleated myofibers than in non-centrally-nucleated myofibers.

## Discussion

This study adds to the growing literature reporting spatial restriction of dystrophin protein to regions of sarcolemma proximal to its myonucleus of origin. Analyses in *mdx52-Xist*^Δhs^ mice revealed two distinct types of spatial phenomena. Firstly, a ‘zebra-like’ patchy pattern of dystrophin was observed in the majority of myofibers, as is expected from the underlying pattern of skewed X-chromosome inactivation of the healthy *Dmd* allele, and consistent with previous observations (**Figures 1,2,4,5,8**).^9,11,14^ An increase in overall dystrophin protein expression levels was observed in aged vs. adult *mdx52-Xist*^Δhs^ animals (**Figures 1C,4C**), consistent with the notion that newly-dystrophin positive myofibers will tend to accumulate over time, as a consequence of a positive selection.^4,41^ Nevertheless, aged *mdx52-Xist*^Δhs^ animals still exhibited non-uniform patterns of sarcolemmal dystrophin, indicating that accumulation of dystrophin-expressing regions is insufficient to completely resolve the observed sarcolemmal patchiness (**Figure 5**). These are disease-relevant observations which mirror situations in which dystrophic myofibers may exist as heterokaryons containing both dystrophin-expressing and non-dystrophin-expressing myonuclei. Specifically, in the case of female dystrophinopthay,^41,42^ and in dystrophic muscle after partially-effective dystrophin restoration strategies. Our group recently showed that CRISPR-Cas9-mediated exon excision restores dystrophin in a patchy manner along the sarcolemma, while PPMO-mediated exon skipping results in a uniform dystrophin distribution.^8–10^ The patchy pattern of dystrophin observed for the former strategy was attributed to productive editing of the *Dmd* gene occurring in only a subset of myonuclei.^10^ As such, CRISPR-Cas9-treated dystrophic muscles are comprised of mosaic myofibers, a situation that is modelled by the *mdx52*-*Xist*^Δhs^ mouse. Subsequently, Morin *et al*., reported similar differential patterns of dystrophin restoration upon CRISPR-Cas9 gene editing or exon-skipping using tri-cyclo-DNA (tcDNA) oligomers.^11^ Importantly, other types of dystrophin-restoration strategy have the potential to generate myofiber heterokaryons, and consequently patchy sarcolemmal dystrophin coverage. For example, in the case of cell therapy, dystrophin-expressing nuclei fuse with otherwise dystrophin negative myofibers.^43–45^

Myofibers expressing dystrophin in a patchy manner are likely to be susceptible to cycles of damage and repair. This notion is supported by the observation that total dystrophin protein expression and the proportion of centrally-nucleated fibers are inversely correlated in *mdx52-Xist*^Δhs^ mice (**Figures 3,6**). This suggests that patchy dystrophin can, at least to some extent, protect against the development of muscle histopathological features. These results are consistent with previous reports in the *mdx*-*Xist*^Δhs^ and *mdx*/*utrn*^−/−^/*Xist*^Δhs^ models, whereby animals expressing low levels of patchy wild-type dystrophin showed higher proportions of centrally-nucleated fibers in comparison to other groups.^14,15^

Concerning the second spatial phenomenon, we observed that dystrophin, and dystrophin-associated proteins, were absent from centrally-nucleated myofibers and myofiber segments (**Figures 7,8,9,S4**). This surprising finding suggests that centrally-nucleated myofibers are refractory to dystrophin expression, at least in this model. Our first thought was that dystrophin may be absent as a consequence of myofiber immaturity.^46^ However, several lines of evidence argue against this notion. Firstly, dystrophin absence in centrally-nucleated regions was observed in 60-week-old mice (**Figure 8,S4**), when regeneration events are likely to be very limited (as also evidenced by limited staining for utrophin, a marker of regenerating myofibers,^47^ **Figure S2**). Secondly, *mdx52-Xist*^Δhs^ centrally-nucleated fibers also exhibit; (i) sizes consistent with healthy (or hypertrophic) myofibers (**Figures 10,11**), (ii) expression of late-stage markers of muscle differentiation like TTN (**Figures 8,9**), and (iii) frequently contained multiple chains of central nuclei (**Figures 8,9,11,S4,S5,S7**). Together, these data suggest that these are in fact mature myofibers, exhibiting signs of repeated historical degeneration and accumulated repair. A second possible explanation is that the absence of dystrophin in centrally-nucleated myofibers and myofiber segments is simply a consequence of the XCI effect, whereby myonuclei that lack the capacity to expresses dystrophin have become clustered in the same region by chance. However, this explanation is not supported by the data, as if this were true, we would expect to observe centrally-nucleated myofibers that exhibit the ‘zebra-like’ patchy pattern of dystrophin throughout, of which we observed none (**Figure 7,S4**). As such, it is unlikely that central nucleation-associated absence of dystrophin expression can be explained by model-associated XCI effects alone.

Myoinjury by BaCl_2_ injection in wild-type mice did not recapitulate the effect observed in *mdx52-Xist*^Δhs^ mice (i.e. post-regeneration, centrally-nucleated myofibers were uniformly dystrophin positive, **Figure S6**). As such, the absence of dystrophin in the centrally-nucleated myofibers of *mdx52-Xist*^Δhs^ mice is likely the result of an interaction between the post-regeneration and dystrophic environments. In further support, the myofibers of X-linked myotubular myopathy patients (and animal models thereof) exhibit centrally-nucleated myofibers and express dystrophin.^48^ Similarly, centronuclear myopathy patients are also known to express dystrophin, albeit with an abnormal intra-cytoplasmic localization.^49^ These observations suggest that central nucleation *per se* is insufficient to prevent dystrophin expression.

*Dmd* mRNA was found to be uniformly distributed throughout *mdx52*-*Xist*^Δhs^ myofibers (both centrally-nucleated and non-centrally-nucleated regions) with all myonuclei containing nuclear ‘blobs’ that are characteristic of *Dmd* RNA-FISH signal (**Figure 9A,B**). These findings suggest that there is no impairment in *Dmd* transcription in these regions. Furthermore, the uniform expression of TTN and F-actin proteins (**Figure 9B,C**) suggests that there is no global impairment in translation for centrally-nucleated regions. We therefore conclude that dystrophin is specifically repressed at the level of translation in *mdx52*-*Xist*^Δhs^ centrally-nucleated myofibers/myofiber segments. Further work is needed to determine the mechanism of this repression, although it is tempting to speculate that local accumulation of *trans*-acting factors, such as miRNAs, may be responsible. For example, miR-31 has been reported to repress dystrophin expression and is upregulated in DMD patient biopsies and *mdx* muscle tissues.^27,50,51^ Likewise, other miRNAs (miR-146a, miR-146b, and miR-374a) have also been reported to repress dystrophin.^52,53^

The absence of dystrophin in centrally-nucleated myofibers/fiber-segments is a disease-relevant observation, as it is suggestive of an additional challenge to the successful re-introduction of dystrophin protein in dystrophic muscle, which may limit the effectiveness of current and future experimental therapeutic interventions. Indeed, such a discrepancy between RNA-level exon skipping levels and dystrophin protein levels after antisense oligonucleotide treatment in DMD patients has been previously reported.^54^ Importantly, we have previously reported widespread rescue of dystrophin expression after antisense oligonucleotide-mediated exon skipping with highly potent PPMO compounds,^8,9^ and similar findings have been reported by others using various dystrophin restoration strategies,^55–59^ which would appear to contradict with the findings reported herein. Notably, treated animals are typically analysed using transverse muscle sections (with isolated single myofiber analysis being relatively rare). As such, some within-fiber patchiness may have been obscured in these analyses. Alternatively, high levels of exon skipping may have been sufficient to overcome the mechanism that represses dystrophin protein expression in centrally-nucleated fibers (i.e. there is a threshold effect).

The microtubule network was found to be disrupted in *mdx52-Xist*^Δhs^ mice at a level intermediate between that of C57 and *mdx52* mice (**Figure 10A-C**). However, microtubule network organization was found to be similar between dystrophin-positive and dystrophin-negative myofiber regions in regions with ‘zebra-like’ patterns of dystrophin expression (**Figure 10D,E**). Conversely, pronounced differences were observed in microtubule organization when comparing centrally-nucleated and non-centrally-nucleated fibers from *mdx52* mice, suggesting that impaired trafficking of dystrophin mRNA and/or protein may contribute to its translational repression in the case of the CNF-associated phenomenon (**Figure 11E-G**). This aligns with previous findings by Percival *et al.* who demonstrated aberrant distribution and increased density of Golgi elements at the surface of centrally nucleated wild-type fibres after cardiotoxin injury (in comparison to non-CNF).^32^

The myonuclear domain theory posits that each nucleus within a syncytial myofiber controls gene expression and protein synthesis within a limited volume of surrounding sarcoplasm.^13^ This concept is helpful for explaining some spatially-restricted gene expression features in syncytial skeletal myofibers. Nevertheless, the definition of myonuclear domain is somewhat flexible (i.e. not restricted to a specific volume or sarcolemmal distance), but to a certain extent abstract and context dependent. Although the size of myonuclear domains can be approximated, it likely varies from cell to cell and protein to protein. Moreover, it is unclear how the myonuclear domain theory would apply to centrally nucleated myofibers, which are not only hypernucleated but also contain chains of seemingly compressed nuclei, that often run in parallel to each other (**Figure 8,S5, S7**).

The *mdx52*-*Xist*^Δhs^ model presents a unique opportunity to study dystrophin-dependent, and CNF-associated spatial phenomena in myofiber heterokaryons. However, it remains to be determined if such effects are present in DMD patient muscle. Dystrophin patchiness has been reported in female dystrophinopathy,^41,42^ and Torelli *et al*., have reported an inverse relationship between differential sarcoplasmic dystrophin coverage and disease severity in patient biopsies.^12^ Notably, patient biopsy material is typically analysed in transverse orientation, whereas dystrophin patchiness is more readily apparent in longitudinal sections. Importantly, centrally-located myonuclei are known to migrate to the myofiber periphery following the completion of regeneration in human muscle, in contrast with the situation in mouse.^33,34^ However, experimentally determining whether central-nucleation-associated impairment in dystrophin expression similarly occurs in human muscle may be challenging. Analyses in healthy or DMD patient muscle would be inadequate, as these either express 100% or close to 0% dystrophin (accounting for a small number of dystrophin-expressing revertant fibers), respectively. The closest analogous situation would be that of female dystrophinopathy, or treated muscle with incomplete dystrophin restoration. Female dystrophinopathy is relatively rare (~2-22% of carrier females),^60,61^ and there is a wide range of pathological presentation and XCI involvement,^62^ which would complicate these analyses. Furthermore, isolated single myofiber analyses in patient muscle are uncommon due to the requirement for fresh material. Myofiber necrosis and regeneration are also known to be more prominent in mouse models than in DMD patients.^63^

In conclusion, this work has identified two spatially-restricted dystrophin expression phenomena within the sarcolemma of a novel dystrophic mouse model. Local expression of dystrophin points to a previously unappreciated level of subcellular complexity in gene expression regulation with important implications for efforts to restore dystrophin protein expression in the muscles of DMD patients.

## Methods

### Animal studies

All experimental procedures were approved by the UK home office, under the project license number PP6777529 (Oxford) or PPL 70/7777 (RVC, approved by the Royal Veterinary College Animal Welfare and Ethical Review Board), in accordance with the Animals (Scientific Procedures) Act 1986. Animals were housed in individually ventilated cages with a 12:12 hour light:dark cycle, with food and water provided *ad libitum*.

*Xist*^Δhs^ animals were a kind gift from Prof. Neil Brockdorff (University of Oxford).^18^ *Xist*^Δhs^ animals contain a deletion of DNase hypersensitivity region upstream of the P1 promoter of the *Xist* gene, resulting in preferential silencing of the mutation-containing chromosome. In heterozygous animals, the mutated X-chromosome is inactivated in up to 90% of the cells.^18^ The *Xist*^Δhs^ mouse has a mixed genetic background consisting of C57BL/6 and CBA.

*mdx52-Xist*^Δhs^ animals were generated by crossing male *mdx52* animals with female *Xist*^Δhs^ mice, with the resulting female F1 progeny used for experimentation.

Dystrophic *mdx52* (C57BL/6J129S-Dmd^tm1Mok^) animals were a kind gift from Dr. Yoshitsugu Aoki (National Centre of Neurology and Psychiatry, Tokyo, Japan). The line was generated by Dr. Motoya Katsuki via targeted replacement of exon 52 in the *Dmd* gene with a neomycin resistance transgene cassette (in the antisense orientation).^16^

Wild-type C57BL/6JOlaHsd (C57BL/6) mice were obtained from Inotiv (London, England) and served as wild-type control animals. Wild-type C57BL/10J mice were used for the BaCl_2_ injury study.

Myoinjury was induced by injection of 1.2% BaCl_2_ (Sigma-Aldrich, MO, USA) in sterile saline (total volume 20 µl) into TA muscles. Injections were performed under anaesthesia using fentanyl/fluanisone (Hypnorm, Vetapharma, Leeds, UK) and midazolam (Hypnovel, Roche, Welwyn Garden City, UK), as described previously^64^. TA muscles were macrodissected 29 days post injury, flash frozen in liquid nitrogen-cooled isopentane, and samples stored at −80°C until ready for analysis.

### Western blot

Dystrophin protein quantification was performed on tibialis anterior (TA) lysates. For protein extraction, 200 TA sections (8 µm thickness) were lysed in modified Radio-Immunoprecipitation Assay (RIPA) buffer (50 mM Tris pH 8, 150 mM NaCl, 1% IGEPAL CA-630, 0.5% sodium deoxycholate, 10% SDS) containing 1× cOmplete proteinase inhibitors (Merck, NJ, USA). Samples were heated for 3 min at 100°C and centrifuged at room temperature for 10 minutes at 15,800 *g*. Protein concentration was measured using Pierce BCA Protein Assay Kit (Thermo Fisher Scientific, MA, USA) according to the manufacturer’s instructions.

20-40 µg of total protein were prepared in NuPAGE LDS sample buffer supplemented with NuPAGE sample reducing agent (both Thermo Fisher Scientific) and denatured for 10 minutes at 75°C. Standards were prepared as a mix of defined different protein ratios (0-75% of wild-type dystrophin protein levels) isolated from positive control, wild-type C57 and negative control, dystrophic (*mdx52*) mouse TA. Linearity of signal was assumed for the few samples that fell outside of the standard range. All samples were loaded onto a pre-cast, NuPAGE Tris-Acetate (3-8%, Thermo Fisher Scientific) and electrophoresis run at 130 V for 1 hour 45 minutes in NuPAGE Tris-Acetate SDS Running Buffer (Thermo Fisher Scientific). Protein was electrotransferred onto 0.45 µm polyvinylidene fluoride (PVDF) membranes (Merck) for 1 hour at 30 V followed by 1 hour at 100 V in 1× NuPAGE Transfer Buffer (Thermo Fisher Scientific) supplemented with 0.1 g/l of SDS (Sigma-Aldrich) and 20% methanol. Total protein was visualized using a ChemiDoc Imaging system (Bio-Rad, CA, USA) measuring fluorescence at 700 nm. The membrane was then washed in wash solution and blocked in blocking solution (either Odyssey blocking buffer (LI-COR Biosciences, NE, USA) or 5% milk (w/v) in tris-buffered saline buffer supplemented with 0.15 Tween-20 (v/v, TBST)). Membranes were incubated with mouse anti-dystrophin, mouse anti-utrophin or mouse anti-vinculin primary antibody (**Table S1**) overnight in blocking buffer at 4°C. Membranes were washed in tris-buffered saline buffer with 0.1% Tween-20 v/v (TBST) and incubated with anti-mouse IgG horseradish peroxidase (HRP) linked antibody (**Table S2**) in blocking buffer + 0.1% Tween-20 for 1 hour at room temperature. Chemiluminescence signal was detected using Clarity Western enhanced chemiluminescence (ECL) substrate (Bio-Rad). If membrane re-probing was necessary for the detection of proteins of similar molecular mass (e.g. dystrophin and utrophin) or using different antibodies from the same host, the membrane was stripped in 0.2 M sodium hydroxide (NaOH) for 30-120 minutes at room temperature. Subsequently, blocking step, primary and secondary antibody incubation and HRP-based detection were performed as described above.

### Single fiber isolation

Extensor digitorum longus (EDL) single myofiber isolation was performed as described previously.^65^ Briefly, EDL muscles were dissected tendon-to-tendon and incubated in 0.2% collagenase II (Worthington, NJ, USA) diluted in filter-sterilized DMEM (Thermo Fisher Scientific, pre-warmed at 37°C) for 45-52 minutes at 37°C. Digestion was stopped by transferring the muscle into a 3.5 cm cell culture dish, containing FluoroBrite DMEM media (Thermo Fisher Scientific) supplemented with 1% Antibiotic-Antimycotic (PSA: Penicillin, Streptomycin and Amphotericin B; Thermo Fisher Scientific) pre-warmed at 37°C. Single myofibers were released from the muscle by gentle flushing using a 200 µl pipette under a stereomicroscope. Freshly isolated myofibers were transferred into a spot plate containing 4% ultrapure paraformaldehyde solution (PFA, Electron Microscopy Sciences, PA, USA) for fixation for 10 minutes at room temperature. Fixed myofibers were washed twice with ultrapure PBS for 5 minutes at room temperature. Immunofluorescence and/or hybridization chain reaction-based RNA *in situ* hybridisation (HCR-RNA-FISH) were performed immediately after fixation and PBS washes.

### Immunofluorescence in tissue sections

Fresh frozen TA muscles were mounted onto corks with Tissue-TEK optimal cutting temperature (OCT) Compound (Sakura, Japan) and cryosectioned (8 µm) in transverse and longitudinal orientations. Samples were stored at −80°C prior to analysis. On the day of staining, slides were air-dried and soaked in phosphate-buffered saline (PBS, Thermo Fisher Scientific) for 10 minutes at room temperature. Sections were blocked in blocking buffer composed of PBS supplemented with 20% foetal calf serum (FCS, Thermo Fisher Scientific) and 20% normal goat serum (NGS, MP Biomedicals, CA, USA) for 2 hours at room temperature. Subsequently, slides were incubated with primary antibodies (listed in **Table S1**) in blocking buffer for 2 hours at room temperature. After washing 3 times with PBS, slides were incubated with secondary fluorescent antibodies (**Table S2**) in PBS or blocking buffer for 1 hour at room temperature in darkness. Slides were then washed 3 times with PBS, incubated with 4C,6-diamidino-2-phenylindole (DAPI) or Hoechst in PBS (1:5,000, Thermo Fisher Scientific), washed with PBS once more and mounted using Dako, Fluorescence Mounting Medium (Agilent Technologies, CA, USA) or SlowFade Diamond Antifade Mountant (Thermo Fisher Scientific).

### Immunofluorescence in isolated fibers

If protein immunodetection and HCR RNA-FISH were performed on the same myofiber sample, staining for protein was carried out first. PFA-fixed myofibers were permeabilized with 1% Triton-X100 (Sigma-Aldrich) for 10 minutes at room temperature followed by a single wash with ultrapure PBS (Thermo Fisher Scientific) for 5 minutes. Subsequently, blocking was performed for 30 minutes at room temperature with blocking buffer containing either 1% bovine serum albumin (BSA, Sigma-Aldrich) diluted in ddH_2_O if only protein immunodetection was performed or 1% ultrapure BSA with RiboLock RNase Inhibitor at 1 U/µl (both Thermo Fisher Scientific) if protein detection was followed by RNA HCR-FISH (see below). Myofibers were then incubated with primary antibodies (**Table S1**) diluted in blocking buffer for 2 hours at room temperature. Thereafter, myofibers were washed 3 times with PBS (Thermo Fisher Scientific) containing 0.1% of Tween-20 (v/v, PBST, Sigma-Aldrich) at room temperature. Subsequently, myofibers were incubated with secondary fluorescent antibodies (**Table S2**) for 2 hours at room temperature in the dark.

If only protein detection was performed, samples were washed 3 times with PBST and incubated with DAPI (Thermo Fisher Scientific) diluted in PBS for at least 2 minutes at room temperature. Myofibers were transferred onto the SuperFrost Plus microslides (VWR) containing 35 µl of SlowFade Diamond Antifade Mountant (Thermo Fisher Scientific) and covered with High Precision Cover Glasses, 1.5 mm thickness (Thorlabs, NJ, USA). Stained myofibers were imaged on the following day and/or stored at −20 °C for repeated imaging.

If HCR-RNA-FISH was subsequently performed, samples were washed once with PBS and fixed in 4% PFA (Electron Microscopy Sciences) at room temperature.

### RNA**-**FISH

RNA was detected using HCR RNA-FISH products purchased from Molecular Instruments (Los Angeles, CA, USA), following the generic sample in solution protocol with modifications. PFA-fixed myofibers were washed twice with PBS (Thermo Fisher Scientific) for 5 minutes at room temperature. Subsequently, samples were incubated with 2× ultrapure saline-sodium citrate (SSC) buffer diluted in ultrapure H_2_O (both Thermo Fisher Scientific) for 5 minutes at room temperature. Then, samples were incubated in pre-warmed (37°C) hybridization buffer (Molecular Instruments) for 30 minutes at 37°C. HCR probe sets were added to fresh, pre-warmed (37°C) hybridization buffer at 1.25 nM/sample. Myofibers were incubated with the probe set solutions at 37°C in humidified conditions for 12-16 hours. Samples were washed with pre-warmed (37°C) wash buffer (Molecular Instruments) five times for 10 minutes at 37°C followed by two washes with 5C SSCT (SSC supplemented with 0.1% v/v Tween-20) at room temperature. Samples were subsequently incubated with amplification buffer (Molecular Instruments) for 30 minutes at room temperature (pre-amplification). Hairpin amplifiers were heated at 95°C for 90 seconds and cooled to room temperature in the dark for 30 minutes. After pre-amplification, samples were incubated with hairpin amplifiers mixed in amplification buffer for 3.5-4 hour at room temperature in darkness. Subsequently, samples were washed 5 times with 5× SSCT for 10 minutes and incubated with 0.1 µg/ml DAPI diluted in PBS (both Thermo Fisher Scientific) for at least 2 min at room temperature. Myofibers were transferred onto the SuperFrost Plus microslides (VWR) and mounted using SlowFade Diamond Antifade Mountant (Thermo Fisher Scientific) and High Precision Cover Glasses (Thorlabs). Stained myofibers were imaged on the following day and/or stored at −20°C for repeated imaging.

### Microscopy

Immunofluorescence microscopy of tissue sections was performed using either wide-field Leica DMIRB Inverted Microscope with MetaMorph imaging software (Molecular Devices, CA, USA) or wide-field Leica DMi8 fluorescence microscope with LAS X Microscope Science Software Platform (all Leica Microsystems, Wetzlar, Germany). For each protein staining, optimal exposure time was chosen based on negative staining control where samples were incubated with secondary antibodies only, to account for background noise and autofluorescence of tissues. All images were processed using Fiji software.^66^ Standard image processing for tissue section images included background subtraction (based on rolling ball with radius of 50 pixels), and brightness and contrast adjustment.

Single myofiber imaging was performed with ZEISS LSM 980 confocal microscope with Airyscan2 detector (ZEISS, Oberkochen, Germany). Depending on the application, the following objectives were used: 40× Plan-Apochromat oil objective (numerical aperture NA = 1.4), 25× Plan-Apochromat (NA = 0.8) or 20× Plan-Apochromat (NA = 0.8). The choice of the objective was based on the field of view and detail required in each experiment.

### Image Analysis

#### CNF and CSA quantification in transverse tissue sections

Immunofluorescence was performed in fresh frozen TA sections using primary antibodies against α2-laminin (**Table S1**) to mark the muscle membrane, and DAPI (Thermo Fisher Scientific) to label nuclei. The proportion of CNFs and myofiber cross-sectional area (CSA) was analysed in transverse TA muscle sections using an open-source Fiji plugin: MuscleJ2, according to developer’s instructions.^67^ 2-5 whole TA sections were analysed per animal. For CNF proportion analysis, values for multiple sections from the same mouse were averaged.

#### Classification of centrally nucleated, segmented and non-centrally nucleated myofibers

Classification into non-centrally nucleated (non-CNF), segmented, and CNFs, and assessment of DAPC protein expression was performed on myofibers isolated from *mdx52-Xist*^Δhs^ mice (aged between 12 and 17 weeks). Myofibers were isolated and stained as described above, using the antibodies listed in **Table S1**. Each slide, was scanned using the wide-field Leica DMi8 fluorescence microscope to visualize all myofibers. Each myofiber was visually examined for nuclei (DAPI), and DAPC protein signal at the sarcolemma and classified according to a pre-defined decision schema (**Figure S8**). Briefly, myofibers were first classified into non-CNF, segmented and CNF groups based on nuclear DAPI staining. Both myofiber classes were further grouped into DAPC-expressing and non-expressing groups. Due to the observation that DAPC is present at the neuromuscular and myotendinous junctions of almost all *mdx52-Xist*^Δhs^ myofibers, junctional sarcolemma regions were excluded from the analysis.

### Analysis of the microtubule network organization

Microtubule intersection angle was analysed using TeDT direction v2017 according to the developer’s instructions.^68^ Two to three cortical microtubule regions per myofiber segment were analysed from z-stack images acquired at 40× magnification. Pre-analysis image processing included the z-projection of cortical microtubule region and background subtraction (radius = 50 pixels). Images were arranged so that transverse microtubules were positioned at a 90° angle with respect to the longitudinal axis of the myofiber. The isotropic areas of microtubule nucleation surrounding myonuclei were excluded from the analysis.^40,68^ In total *n* = 60 regions from 32 myofibers, 79 regions from 40 myofibers, and 65 regions from 31 myofibers were analysed for C57, *mdx52-Xist*^Δhs^, and *mdx52* mice respectively. The histogram of proportion of microtubule directionality angles for each genotype was prepared using averaged and 0-1 normalized values of fractions of microtubules at 0-176° in 4° intervals.

### Serum microRNA analysis

Serum miRNA analysis was performed as described previously.^69,70^ Briefly, RNA was extracted from 50 µl blood serum samples using TRIzol LS (Thermo Fisher Scientific) according to manufacturer’s instructions with minor modifications. A synthetic spike-in control oligonucleotide with a non-mammalian miRNA sequence (i.e. cel-miR-39, 2.5 fmol, IDT) was added at the phenolic extraction phase in order to allow for between-sample normalization. miRNAs were quantified using the small RNA TaqMan RT-qPCR method using miRNA-specific stem loop reverse transcription primers. Details of miRNA assays are listed in **Table S3**. Reverse transcription was performed using the TaqMan MicroRNA Reverse Transcription Kit (Thermo Fisher Scientific). cDNA was amplified using a StepOne Plus real-time PCR thermocycler with TaqMan Gene Expression Master Mix (both Thermo Fisher Scientific) using universal cycling conditions: 95°C for 10 minutes, followed by 40 cycles of 95°C for 15 seconds and 60°C for 1 minute. All samples were analyzed in duplicate. Relative quantification was performed using the Pfaffl method,^71^ and miRNA-of-interest abundance normalized to cel-miR-39.^70^

### Statistical Analysis

Statistical analyses were performed using GraphPad Prism (v10.2.3) (GraphPad Software Inc., San Diego, California, USA). For comparisons of two groups, a Student’s *t*-test was used. For comparisons of more than two groups, an ordinary one-way analysis of variance (ANOVA) was performed with Bonferroni’s *post hoc* test for inter-group comparisons

## Acknowledgements

KC was supported by doctoral studentship from the Clarendon Fund in partnership with the Medical Research Council (MRC), and the Juel-Jenson Scholarship from St Cross College, Oxford. This work was supported by a grant from the UK Medical Research Council (awarded to MJAW and TCR).

## Author Contributions

TCR, KC, and MJAW conceived the study. TCR, RP, ETW, and MJAW supervised the work. KC, VF, NH, JCWH, and LER performed experimentation. AAR and MvP provided essential reagents. TCR wrote the first draft of the manuscript. All authors contributed to the final version of the manuscript.

## Declaration of Interests

MJAW discloses being an advisor and shareholder in PepGen Ltd, a biotechnology company that aims to generate exon skipping therapies for DMD. MJAW has filed multiple patents relating to exon skipping technologies for treating DMD. AAR discloses being employed by LUMC which has patents on exon skipping technology, some of which has been licensed to BioMarin and subsequently sublicensed to Sarepta. As co-inventor of some of these patents AAR was entitled to a share of royalties. AAR further discloses being ad hoc consultant for PTC Therapeutics, Sarepta Therapeutics, Regenxbio, Dyne Therapeutics, Lilly, BioMarin Pharmaceuticals Inc., Eisai, Entrada, Takeda, Splicesense, Galapagos, Sapreme, Italfarmaco and Astra Zeneca. AAR also reports being a member of the scientific advisory boards of Hybridize Therapeutics (past), Silence Therapeutics, Sarepta therapeutics, Sapreme and Mitorx. Remuneration for consulting and advising activities is paid to LUMC. In the past 5 years, LUMC also received speaker honoraria from Alnylam Netherlands, Italfarmaco and Pfizer and funding for contract research from Sapreme, Eisai, BioMarin, Galapagos and Synaffix. Project funding is received from Sarepta Therapeutics and Entrada via unrestricted grants. RJP has received funding for separate research programmes from Pfizer, Ultragenyx, and Exonics Therapeutics and has been a consultant to Exonics Therapeutics; the financial interests were reviewed and approved by the University in accordance with conflict of interest policies. The remaining authors declare no competing financial interests.

## Data Availability Statement

All data are included in the manuscript. Raw data are available on request.

## References

1. Petrof, B.J., Shrager, J.B., Stedman, H.H., Kelly, A.M., and Sweeney, H.L. (1993). Dystrophin protects the sarcolemma from stresses developed during muscle contraction. Proc. Natl. Acad. Sci. U.S.A. 90, 3710–3714.

2. Roberts, T.C., Wood, M.J.A., and Davies, K.E. (2023). Therapeutic approaches for Duchenne muscular dystrophy. Nat Rev Drug Discov, 1–18. 10.1038/s41573-023-00775-6.

3. Amoasii, L., Long, C., Li, H., Mireault, A.A., Shelton, J.M., Sanchez-Ortiz, E., McAnally, J.R., Bhattacharyya, S., Schmidt, F., Grimm, D., et al. (2017). Single-cut genome editing restores dystrophin expression in a new mouse model of muscular dystrophy. Sci Transl Med 9. 10.1126/scitranslmed.aan8081.

4. Hanson, B., Wood, M.J.A., and Roberts, T.C. (2021). Molecular correction of Duchenne muscular dystrophy by splice modulation and gene editing. RNA Biol 18, 1048–1062. 10.1080/15476286.2021.1874161.

5. Nelson, C.E., Hakim, C.H., Ousterout, D.G., Thakore, P.I., Moreb, E.A., Castellanos Rivera, R.M., Madhavan, S., Pan, X., Ran, F.A., Yan, W.X., et al. (2016). In vivo genome editing improves muscle function in a mouse model of Duchenne muscular dystrophy. Science 351, 403–407. 10.1126/science.aad5143.

6. Tabebordbar, M., Zhu, K., Cheng, J.K.W., Chew, W.L., Widrick, J.J., Yan, W.X., Maesner, C., Wu, E.Y., Xiao, R., Ran, F.A., et al. (2016). In vivo gene editing in dystrophic mouse muscle and muscle stem cells. Science 351, 407–411. 10.1126/science.aad5177.

7. Muntoni, F., Maresh, K., Davies, K., Harriman, S., Layton, G., Rosskamp, R., Russell, A., Tejura, B., and Tinsley, J. (2017). PhaseOut DMD: a Phase 2, proof of concept, clinical study of utrophin modulation with ezutromid. Neuromuscular Disorders 27, S217. 10.1016/j.nmd.2017.06.443.

8. Chwalenia, K., Oieni, J., Zemła, J., Lekka, M., Ahlskog, N., Coenen-Stass, A.M.L., McClorey, G., Wood, M.J.A., Lomonosova, Y., and Roberts, T.C. (2022). Exon skipping induces uniform dystrophin rescue with dose-dependent restoration of serum miRNA biomarkers and muscle biophysical properties. Molecular Therapy - Nucleic Acids 29, 955–968. 10.1016/j.omtn.2022.08.033.

9. van Westering, T.L.E., Lomonosova, Y., CoenenCStass, A.M.L., Betts, C.A., Bhomra, A., Hulsker, M., Clark, L.E., McClorey, G., AartsmaCRus, A., van Putten, M., et al. (2020). Uniform sarcolemmal dystrophin expression is required to prevent extracellular microRNA release and improve dystrophic pathology. Journal of Cachexia, Sarcopenia and Muscle 11, 578–593. 10.1002/jcsm.12506.

10. Hanson, B., Stenler, S., Ahlskog, N., Chwalenia, K., Svrzikapa, N., Coenen-Stass, A.M.L., Weinberg, M.S., Wood, M.J.A., and Roberts, T.C. (2022). Non-uniform dystrophin re-expression after CRISPR-mediated exon excision in the dystrophin/utrophin double-knockout mouse model of DMD. Molecular Therapy - Nucleic Acids 30, 379–397. 10.1016/j.omtn.2022.10.010.

11. Morin, A., Stantzou, A., Petrova, O.N., Hildyard, J., Tensorer, T., Matouk, M., Petkova, M.V., Richard, I., Manoliu, T., Goyenvalle, A., et al. (2023). Dystrophin myonuclear domain restoration governs treatment efficacy in dystrophic muscle. Proceedings of the National Academy of Sciences 120, e2206324120. 10.1073/pnas.2206324120.

12. Torelli, S., Scaglioni, D., Sardone, V., Ellis, M.J., Domingos, J., Jones, A., Feng, L., Chambers, D., Eastwood, D.M., Leturcq, F., et al. (2021). High-Throughput Digital Image Analysis Reveals Distinct Patterns of Dystrophin Expression in Dystrophinopathy Patients. J Neuropathol Exp Neurol 80, 955–965. 10.1093/jnen/nlab088.

13. Pavlath, G.K., Rich, K., Webster, S.G., and Blau, H.M. (1989). Localization of muscle gene products in nuclear domains. Nature 337, 570–573. 10.1038/337570a0.

14. van Putten, M., Hulsker, M., Nadarajah, V.D., van Heiningen, S.H., van Huizen, E., van Iterson, M., Admiraal, P., Messemaker, T., den Dunnen, J.T., ’t Hoen, P.A.C., et al. (2012). The effects of low levels of dystrophin on mouse muscle function and pathology. PLoS ONE 7, e31937. 10.1371/journal.pone.0031937.

15. van Putten, M., Hulsker, M., Young, C., Nadarajah, V.D., Heemskerk, H., van der Weerd, L., ’t Hoen, P.A.C., van Ommen, G.-J.B., and Aartsma-Rus, A.M. (2013). Low dystrophin levels increase survival and improve muscle pathology and function in dystrophin/utrophin double-knockout mice. FASEB J. 27, 2484–2495. 10.1096/fj.12-224170.

16. Araki, E., Nakamura, K., Nakao, K., Kameya, S., Kobayashi, O., Nonaka, I., Kobayashi, T., and Katsuki, M. (1997). Targeted disruption of exon 52 in the mouse dystrophin gene induced muscle degeneration similar to that observed in Duchenne muscular dystrophy. Biochem Biophys Res Commun 238, 492–497. 10.1006/bbrc.1997.7328.

17. van Westering, T.L.E., Johansson, H.J., Hanson, B., Coenen-Stass, A.M.L., Lomonosova, Y., Tanihata, J., Motohashi, N., Yokota, T., Takeda, S., Lehtiö, J., et al. (2020). Mutation-independent Proteomic Signatures of Pathological Progression in Murine Models of Duchenne Muscular Dystrophy. Mol Cell Proteomics 19, 2047–2067. 10.1074/mcp.RA120.002345.

18. Newall, A.E., Duthie, S., Formstone, E., Nesterova, T., Alexiou, M., Johnston, C., Caparros, M.L., and Brockdorff, N. (2001). Primary non-random X inactivation associated with disruption of Xist promoter regulation. Hum. Mol. Genet. 10, 581–589.

19. Viggiano, E., Ergoli, M., Picillo, E., and Politano, L. (2016). Determining the role of skewed X-chromosome inactivation in developing muscle symptoms in carriers of Duchenne muscular dystrophy. Hum Genet 135, 685–698. 10.1007/s00439-016-1666-6.

20. Sun, M.-X., Jing, M., Hua, Y., Wang, J.-B., Wang, S.-Q., Chen, L.-L., Ju, L., and Liu, Y.-S. (2024). A female patient carrying a novel DMD mutation with non-random X-chromosome inactivation from a DMD family. BMC Medical Genomics 17, 46. 10.1186/s12920-024-01794-x.

21. Hakim, C.H., Wasala, N.B., Nelson, C.E., Wasala, L.P., Yue, Y., Louderman, J.A., Lessa, T.B., Dai, A., Zhang, K., Jenkins, G.J., et al. AAV CRISPR editing rescues cardiac and muscle function for 18 months in dystrophic mice. JCI Insight 3, e124297. 10.1172/jci.insight.124297.

22. Mizuno, Y., Nonaka, I., Hirai, S., and Ozawa, E. (1993). Reciprocal expression of dystrophin and utrophin in muscles of Duchenne muscular dystrophy patients, female DMD-carriers and control subjects. J. Neurol. Sci. 119, 43–52.

23. Culle, M.J., Walsh, J.M., Tinsle, J.M., Fisher, R., and Davies, K.E. (2001). Immunogold confirmation that utrophin is localized to the normal position of dystrophin in dystrophin-negative transgenic mouse muscle. Histochem J 33, 579–583. 10.1023/a:1014964127156.

24. Coenen-Stass, A.M.L., Wood, M.J.A., and Roberts, T.C. (2017). Biomarker Potential of Extracellular miRNAs in Duchenne Muscular Dystrophy. Trends Mol Med 23, 989– 1001. 10.1016/j.molmed.2017.09.002.

25. Roberts, T.C., Godfrey, C., McClorey, G., Vader, P., Briggs, D., Gardiner, C., Aoki, Y., Sargent, I., Morgan, J.E., and Wood, M.J.A. (2013). Extracellular microRNAs are dynamic non-vesicular biomarkers of muscle turnover. Nucl. Acids Res. 41, 9500–9513. 10.1093/nar/gkt724.

26. Coenen-Stass, A.M.L., Betts, C.A., Lee, Y.F., Mäger, I., Turunen, M.P., El Andaloussi, S., Morgan, J.E., Wood, M.J.A., and Roberts, T.C. (2016). Selective release of muscle-specific, extracellular microRNAs during myogenic differentiation. Hum. Mol. Genet. 25, 3960–3974. 10.1093/hmg/ddw237.

27. Coenen-Stass, A.M.L., Sork, H., Gatto, S., Godfrey, C., Bhomra, A., Krjutškov, K., Hart, J.R., Westholm, J.O., O’Donovan, L., Roos, A., et al. (2018). Comprehensive RNA-Sequencing Analysis in Serum and Muscle Reveals Novel Small RNA Signatures with Biomarker Potential for DMD. Mol Ther Nucleic Acids 13, 1–15. 10.1016/j.omtn.2018.08.005.

28. Denes, L.T., Kelley, C.P., and Wang, E.T. (2021). Microtubule-based transport is essential to distribute RNA and nascent protein in skeletal muscle. Nat Commun 12, 6079. 10.1038/s41467-021-26383-9.

29. Scarborough, E.A., Uchida, K., Vogel, M., Erlitzki, N., Iyer, M., Phyo, S.A., Bogush, A., Kehat, I., and Prosser, B.L. (2021). Microtubules orchestrate local translation to enable cardiac growth. Nat Commun 12, 1547. 10.1038/s41467-021-21685-4.

30. Prins, K.W., Humston, J.L., Mehta, A., Tate, V., Ralston, E., and Ervasti, J.M. (2009). Dystrophin is a microtubule-associated protein. J Cell Biol 186, 363–369. 10.1083/jcb.200905048.

31. Belanto, J.J., Mader, T.L., Eckhoff, M.D., Strandjord, D.M., Banks, G.B., Gardner, M.K., Lowe, D.A., and Ervasti, J.M. (2014). Microtubule binding distinguishes dystrophin from utrophin. Proc Natl Acad Sci U S A 111, 5723–5728. 10.1073/pnas.1323842111.

32. Percival, J.M., Gregorevic, P., Odom, G.L., Banks, G.B., Chamberlain, J.S., and Froehner, S.C. (2007). rAAV6-microdystrophin rescues aberrant Golgi complex organization in mdx skeletal muscles. Traffic 8, 1424–1439. 10.1111/j.1600-0854.2007.00622.x.

33. Collins, B.C., Shapiro, J.B., Scheib, M.M., Musci, R.V., Verma, M., and Kardon, G. (2024). Three-dimensional imaging studies in mice identify cellular dynamics of skeletal muscle regeneration. Dev Cell 59, 1457–1474.e5. 10.1016/j.devcel.2024.03.017.

34. Meyer, G.A. (2018). Evidence of induced muscle regeneration persists for years in the mouse. Muscle Nerve 58, 858–862. 10.1002/mus.26329.

35. Guiraud, S., Edwards, B., Squire, S.E., Moir, L., Berg, A., Babbs, A., Ramadan, N., Wood, M.J., and Davies, K.E. (2019). Embryonic myosin is a regeneration marker to monitor utrophin-based therapies for DMD. Hum Mol Genet 28, 307–319. 10.1093/hmg/ddy353.

36. Massopust, R.T., Lee, Y.I., Pritchard, A.L., Nguyen, V.-K.M., McCreedy, D.A., and Thompson, W.J. (2020). Lifetime analysis of mdx skeletal muscle reveals a progressive pathology that leads to myofiber loss. Sci Rep 10, 17248. 10.1038/s41598-020-74192-9.

37. Duddy, W., Duguez, S., Johnston, H., Cohen, T.V., Phadke, A., Gordish-Dressman, H., Nagaraju, K., Gnocchi, V., Low, S., and Partridge, T. (2015). Muscular dystrophy in the mdx mouse is a severe myopathy compounded by hypotrophy, hypertrophy and hyperplasia. Skeletal Muscle 5, 16. 10.1186/s13395-015-0041-y.

38. Echigoya, Y., Lee, J., Rodrigues, M., Nagata, T., Tanihata, J., Nozohourmehrabad, A., Panesar, D., Miskew, B., Aoki, Y., and Yokota, T. (2013). Mutation types and aging differently affect revertant fiber expansion in dystrophic mdx and mdx52 mice. PLoS One 8, e69194. 10.1371/journal.pone.0069194.

39. Bugnard, E., Zaal, K.J.M., and Ralston, E. (2005). Reorganization of microtubule nucleation during muscle differentiation. Cell Motil Cytoskeleton 60, 1–13. 10.1002/cm.20042.

40. Oddoux, S., Zaal, K.J., Tate, V., Kenea, A., Nandkeolyar, S.A., Reid, E., Liu, W., and Ralston, E. (2013). Microtubules that form the stationary lattice of muscle fibers are dynamic and nucleated at Golgi elements. J Cell Biol 203, 205–213. 10.1083/jcb.201304063.

41. Pegoraro, E., Schimke, R.N., Garcia, C., Stern, H., Cadaldini, M., Angelini, C., Barbosa, E., Carroll, J., Marks, W.A., Neville, H.E., et al. (1995). Genetic and biochemical normalization in female carriers of Duchenne muscular dystrophy: evidence for failure of dystrophin production in dystrophin-competent myonuclei. Neurology 45, 677–690.

42. Schmidt-Achert, M., Fischer, P., Müller-Felber, W., Mudra, H., and Pongratz, D. (1993). Heterozygotic gene expression in endomyocardial biopsies: a new diagnostic tool confirms the Duchenne carrier status. Clin Investig 71, 247–253.

43. Gussoni, E., Blau, H.M., and Kunkel, L.M. (1997). The fate of individual myoblasts after transplantation into muscles of DMD patients. Nat Med 3, 970–977. 10.1038/nm0997-970.

44. Partridge, T.A., Morgan, J.E., Coulton, G.R., Hoffman, E.P., and Kunkel, L.M. (1989). Conversion of mdx myofibres from dystrophin-negative to -positive by injection of normal myoblasts. Nature 337, 176–179. 10.1038/337176a0.

45. Gussoni, E., Pavlath, G.K., Lanctot, A.M., Sharma, K.R., Miller, R.G., Steinman, L., and Blau, H.M. (1992). Normal dystrophin transcripts detected in Duchenne muscular dystrophy patients after myoblast transplantation. Nature 356, 435–438. 10.1038/356435a0.

46. Yoshimoto, Y., Ikemoto-Uezumi, M., Hitachi, K., Fukada, S., and Uezumi, A. (2020). Methods for Accurate Assessment of Myofiber Maturity During Skeletal Muscle Regeneration. Front. Cell Dev. Biol. 8. 10.3389/fcell.2020.00267.

47. Lin, S., Gaschen, F., and Burgunder, J.M. (1998). Utrophin is a regeneration-associated protein transiently present at the sarcolemma of regenerating skeletal muscle fibers in dystrophin-deficient hypertrophic feline muscular dystrophy. J Neuropathol Exp Neurol 57, 780–790. 10.1097/00005072-199808000-00007.

48. Lawlor, M.W., Beggs, A.H., Buj-Bello, A., Childers, M.K., Dowling, J.J., James, E.S., Meng, H., Moore, S.A., Prasad, S., Schoser, B., et al. (2016). Skeletal Muscle Pathology in X-Linked Myotubular Myopathy: Review With Cross-Species Comparisons. J Neuropathol Exp Neurol 75, 102–110. 10.1093/jnen/nlv020.

49. Mora, M., Morandi, L., Merlini, L., Vita, G., Baradello, A., Barresi, R., Di Blasi, C., Blasevich, F., Gebbia, M., and Daniel, S. (1994). Fetus-like dystrophin expression and other cytoskeletal protein abnormalities in centronuclear myopathies. Muscle Nerve 17, 1176–1184. 10.1002/mus.880171008.

50. Cacchiarelli, D., Incitti, T., Martone, J., Cesana, M., Cazzella, V., Santini, T., Sthandier, O., and Bozzoni, I. (2011). miR-31 modulates dystrophin expression: new implications for Duchenne muscular dystrophy therapy. EMBO Rep. 12, 136–141. 10.1038/embor.2010.208.

51. Roberts, T.C., Blomberg, K.E.M., McClorey, G., Andaloussi, S.E., Godfrey, C., Betts, C., Coursindel, T., Gait, M.J., Smith, C.E., and Wood, M.J. (2012). Expression Analysis in Multiple Muscle Groups and Serum Reveals Complexity in the MicroRNA Transcriptome of the mdx Mouse with Implications for Therapy. Molecular Therapy — Nucleic Acids 1, e39. 10.1038/mtna.2012.26.

52. Fiorillo, A.A., Heier, C.R., Novak, J.S., Tully, C.B., Brown, K.J., Uaesoontrachoon, K., Vila, M.C., Ngheim, P.P., Bello, L., Kornegay, J.N., et al. (2015). TNF-α-Induced microRNAs Control Dystrophin Expression in Becker Muscular Dystrophy. Cell Rep 12, 1678–1690. 10.1016/j.celrep.2015.07.066.

53. McCormack, N.M., Calabrese, K.A., Sun, C.M., Tully, C.B., Heier, C.R., and Fiorillo, A.A. (2023). Deletion of miR-146a enhances therapeutic protein restoration in model of dystrophin exon skipping. bioRxiv, 2023.05.09.540042. 10.1101/2023.05.09.540042.

54. Inc, W.L.S.U. (2022). Wave Life Sciences Provides Positive Update on Proof-of-Concept Study for WVE-N531 in Duchenne Muscular Dystrophy. GlobeNewswire News Room. https://www.globenewswire.com/news-release/2022/12/19/2576214/0/en/Wave-Life-Sciences-Provides-Positive-Update-on-Proof-of-Concept-Study-for-WVE-N531-in-Duchenne-Muscular-Dystrophy.html.

55. Skuk, D., Goulet, M., Roy, B., Chapdelaine, P., Bouchard, J.-P., Roy, R., Dugré, F.J., Sylvain, M., Lachance, J.-G., Deschênes, L., et al. (2006). Dystrophin expression in muscles of duchenne muscular dystrophy patients after high-density injections of normal myogenic cells. J Neuropathol Exp Neurol 65, 371–386. 10.1097/01.jnen.0000218443.45782.81.

56. Novak, J.S., Hogarth, M.W., Boehler, J.F., Nearing, M., Vila, M.C., Heredia, R., Fiorillo, A.A., Zhang, A., Hathout, Y., Hoffman, E.P., et al. (2017). Myoblasts and macrophages are required for therapeutic morpholino antisense oligonucleotide delivery to dystrophic muscle. Nat Commun 8, 941. 10.1038/s41467-017-00924-7.

57. Cohen, S.A., Bar-Am, O., Fuoco, C., Saar, G., Gargioli, C., and Seliktar, D. (2022). In vivo restoration of dystrophin expression in mdx mice using intra-muscular and intra-arterial injections of hydrogel microsphere carriers of exon skipping antisense oligonucleotides. Cell Death Dis 13, 1–13. 10.1038/s41419-022-05166-0.

58. Liu M, FAU-Yue, Y., Yue Y, FAU-Harper, S.Q., Harper SQ, FAU-Grange, R.W., Grange RW, FAU-Chamberlain, J.S., Chamberlain JS, FAU-Duan, D., et al. Adeno-associated virus-mediated microdystrophin expression protects young mdx muscle from contraction-induced injury. - Mol Ther.2005 Feb;11(2):245–56.

59. Egorova, T.V., Polikarpova, A.V., Vassilieva, S.G., Dzhenkova, M.A., Savchenko, I.M., Velyaev, O.A., Shmidt, A.A., Soldatov, V.O., Pokrovskii, M.V., Deykin, A.V., et al. (2023). CRISPR-Cas9 correction in the DMD mouse model is accompanied by upregulation of Dp71f protein. Mol Ther Methods Clin Dev 30, 161–180. 10.1016/j.omtm.2023.06.006.

60. Hoogerwaard, E.M., Bakker, E., Ippel, P.F., Oosterwijk, J.C., Majoor-Krakauer, D.F., Leschot, N.J., Van Essen, A.J., Brunner, H.G., van der Wouw, P.A., Wilde, A.A., et al. (1999). Signs and symptoms of Duchenne muscular dystrophy and Becker muscular dystrophy among carriers in The Netherlands: a cohort study. Lancet 353, 2116–2119. 10.1016/s0140-6736(98)10028-4.

61. Hoffman, E.P., Arahata, K., Minetti, C., Bonilla, E., and Rowland, L.P. (1992). Dystrophinopathy in isolated cases of myopathy in females. Neurology 42, 967–975. 10.1212/wnl.42.5.967.

62. Finsterer, J., and Stollberger, C. (2018). Muscle, cardiac, and cerebral manifestations in female carriers of dystrophin variants. J Neurol Sci 388, 107–108. 10.1016/j.jns.2018.03.015.

63. Coulton, G.R., Morgan, J.E., Partridge, T.A., and Sloper, J.C. (1988). The mdx mouse skeletal muscle myopathy: I. A histological, morphometric and biochemical investigation. Neuropathol. Appl. Neurobiol. 14, 53–70.

64. Verhaart, I.E.C., Cappellari, O., Tanganyika-de Winter, C.L., Plomp, J.J., Nnorom, S., Wells, K.E., Hildyard, J.C.W., Bull, D., Aartsma-Rus, A., and Wells, D.J. Simvastatin Treatment Does Not Ameliorate Muscle Pathophysiology in a Mouse Model for Duchenne Muscular Dystrophy. J Neuromuscul Dis 8, 845–863. 10.3233/JND-200524.

65. Keire, P., Shearer, A., Shefer, G., and Yablonka-Reuveni, Z. (2013). Isolation and culture of skeletal muscle myofibers as a means to analyze satellite cells. Methods Mol Biol 946, 431–468. 10.1007/978-1-62703-128-8_28.

66. Schindelin, J., Arganda-Carreras, I., Frise, E., Kaynig, V., Longair, M., Pietzsch, T., Preibisch, S., Rueden, C., Saalfeld, S., Schmid, B., et al. (2012). Fiji: an open-source platform for biological-image analysis. Nat Methods 9, 676–682. 10.1038/nmeth.2019.

67. Danckaert, A., Trignol, A., Le Loher, G., Loubens, S., Staels, B., Duez, H., Shorte, S.L., and Mayeuf-Louchart, A. (2023). MuscleJ2: a rebuilding of MuscleJ with new features for high-content analysis of skeletal muscle immunofluorescence slides. Skelet Muscle 13, 14. 10.1186/s13395-023-00323-1.

68. Liu, W., and Ralston, E. (2014). A new directionality tool for assessing microtubule pattern alterations. Cytoskeleton (Hoboken) 71, 230–240. 10.1002/cm.21166.

69. Roberts, T.C., Coenen-Stass, A.M.L., Betts, C.A., and Wood, M.J.A. (2014). Detection and quantification of extracellular microRNAs in murine biofluids. Biol Proced Online 16, 5. 10.1186/1480-9222-16-5.

70. Roberts, T.C., Coenen-Stass, A.M.L., and Wood, M.J.A. (2014). Assessment of RT-qPCR normalization strategies for accurate quantification of extracellular microRNAs in murine serum. PLoS ONE 9, e89237. 10.1371/journal.pone.0089237.

71. Pfaffl, M.W. (2001). A new mathematical model for relative quantification in real-time RT-PCR. Nucleic Acids Res. 29, e45.

