## Supplementary Information for "Myonuclear domain-associated and central nucleation-dependent spatial restriction of dystrophin protein expression in a novel DMD mouse model"

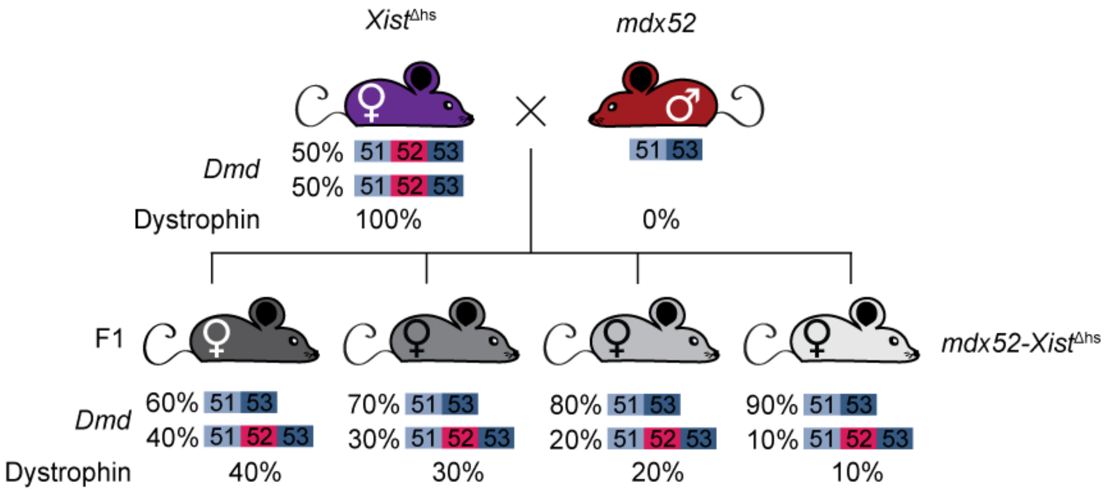


**Figure S1**

**Breeding scheme for *mdx52-Xist*$^{\Delta hs}$ mice.**

Breeding scheme to generate the *mdx52-Xist*$^{\Delta hs}$ model. Female *Xist*$^{\Delta hs}$ mice (containing a deletion in a DNase I hypersensitivity site in the *Xist* promoter) were crossed with male *mdx52* animals (lacking exon 52 of the *Dmd* gene). The resulting female F1 generation (*mdx52-Xist*$^{\Delta hs}$) are expected to exhibit variable levels of dystrophin expression as a consequence of skewed X-chromosome inactivation of the healthy *Dmd* allele. Examples of hypothetical dystrophin expression outcomes are shown (i.e. silencing of the healthy *Dmd*-containing chromosome in 60-90% of cases). For each mouse, the expected X-chromosome inactivation proportions and dystrophin protein expression are indicated.

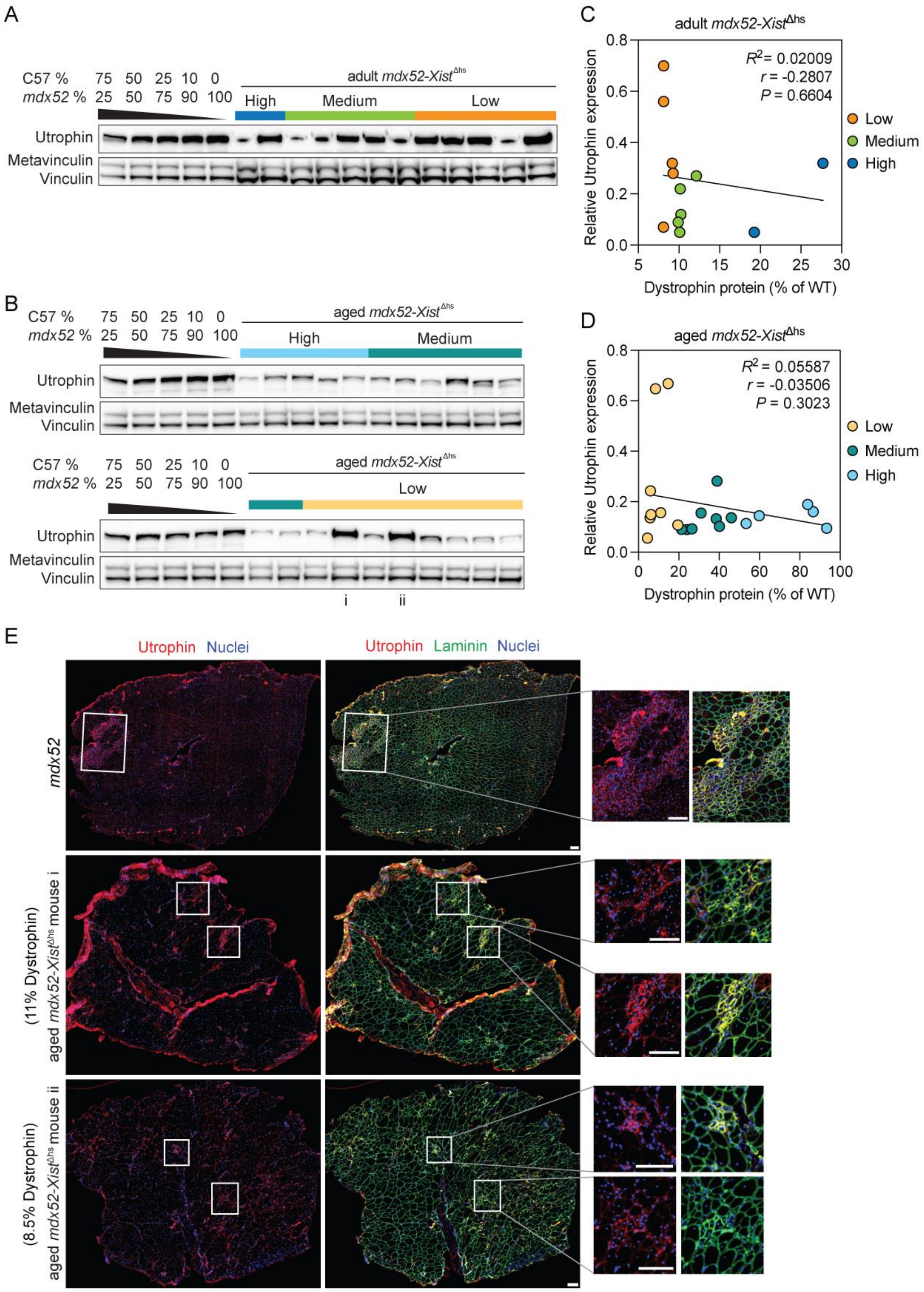
A
C57 %
mdx52 %
75 50 25 10 0
25 50 75 90 100
adult mdx52-Xist^Δhs
High
Medium
Low
Utrophin
Metavinculin
Vinculin
B
aged mdx52-Xist^Δhs
High
Medium
Low
i
ii
C
adult mdx52-Xist^Δhs
R² = 0.02009
r = -0.2807
P = 0.6604
Relative Utrophin expression
Dystrophin protein (% of WT)
D
aged mdx52-Xist^Δhs
R² = 0.05587
r = -0.03506
P = 0.3023
E
Utrophin Nuclei
Utrophin Laminin Nuclei
mdx52
(11% Dystrophin) aged mdx52-Xist^Δhs mouse i
(8.5% Dystrophin) aged mdx52-Xist^Δhs mouse ii

**Figure S2**

**Utrophin expression is associated with muscle regeneration and is not reciprocal with dystrophin expression.**

Western blot analysis of utrophin protein in TA muscle from (**A**) adult (6 weeks old) and (**B**) aged (60 weeks old) *mdx52-Xist*$^{\Delta hs}$ mice. Vinculin was utilised as a loading control. Labelled, coloured boxes demonstrate levels of dystrophin expressed in respective samples. Spearman correlation analysis of dystrophin percentage and relative utrophin abundance in TA muscles of (**C**) adult and (**D**) aged *mdx52-Xist*$^{\Delta hs}$ animals. (**E**) Immunofluorescence staining of utrophin and laminin in TA muscle sections of 12-week-old *mdx52* mouse and 60-week-old *mdx52-Xist*$^{\Delta hs}$ animals from low dystrophin group expressing high levels of utrophin as quantified by western blot. Specific mice with high utrophin expression were selected for immunofluorescence analysis and are labelled as ‘i’ and ‘ii’ on the western blot. Insets demonstrate utrophin signal detected in small-calibre centrally nucleated myofibers in control *mdx52* and both *mdx52-Xist*$^{\Delta hs}$ animals. Values are mean±SD, *n*=2-8. Scale bars represent 100 µm, tiled images taken at 20× magnification and stitched together using LAS X.

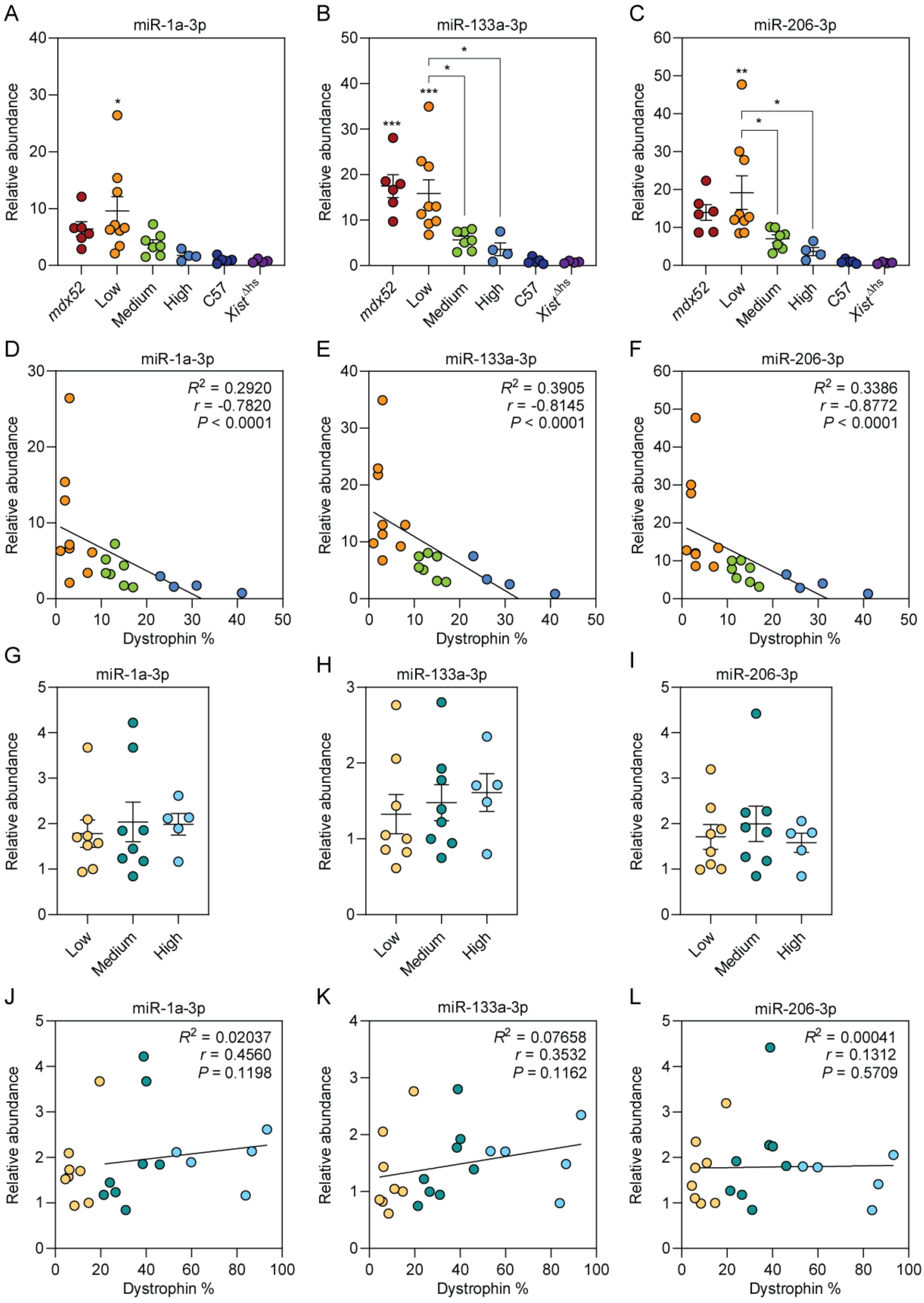
A
miR-1a-3p
Relative abundance
mdx52
Low
Medium
High
C57
Xist^Δhs
B
miR-133a-3p
C
miR-206-3p
D
R² = 0.2920
r = -0.7820
P < 0.0001
Dystrophin %
E
R² = 0.3905
r = -0.8145
P < 0.0001
F
R² = 0.3386
r = -0.8772
P < 0.0001
G
H
I
J
R² = 0.02037
r = 0.4560
P = 0.1198
K
R² = 0.07658
r = 0.3532
P = 0.1162
L
R² = 0.00041
r = 0.1312
P = 0.5709

**Figure S3**

**Analysis of myomiR biomarkers in adult and aged *mdx52-Xist*$^{\Delta hs}$ serum.**

Serum myomiR levels were analysed in adult (6-week-old) *mdx52-Xist*$^{\Delta hs}$ mice and compared with age and sex-matched dystrophic *mdx52*, and wild-type C57 and *Xist*$^{\Delta hs}$ controls for (**A**) miR-1a-3p, (**B**), miR-133a-3p, and (**C**) miR-206-3p. For each miRNA, the relationship between serum abundance and dystrophin protein expression in TA was analysed by Spearman correlation and linear regression (**D**-**F**). Serum myomiR levels were also analysed in aged (60-week-old) *mdx52-Xist*$^{\Delta hs}$ mice for (**G**) miR-1a-3p, (**H**), miR-133a-3p, and (**I**) miR-206-3p, and correlated with dystrophin protein expression in TA as above (**J**-**L**) Statistical significance was assessed by one-way ANOVA and Bonferroni *post hoc* test. Values are mean±SEM, *n*=4-9. **P*<0.05, ***P*<0.01, ****P*<0.001. Statistical comparisons are to the C57 wild-type group unless otherwise indicated.

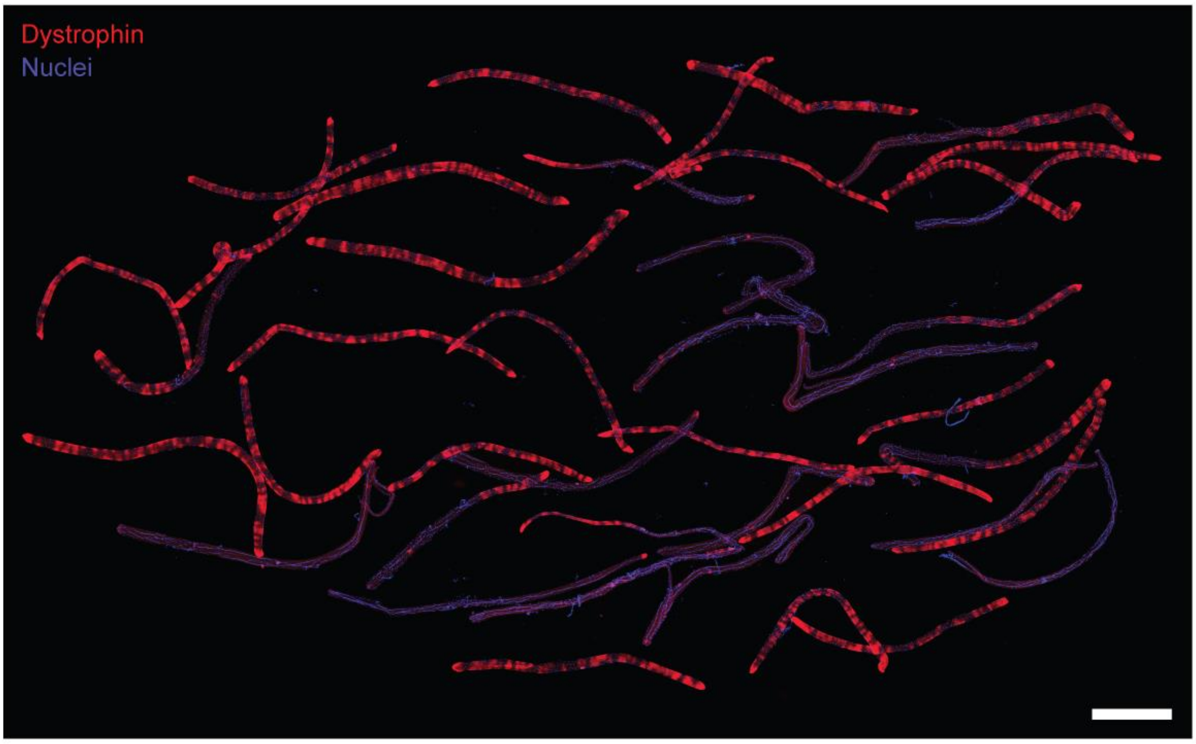


**Figure S4**

**Bulk *mdx52-Xist*$^{\Delta hs}$ myofiber preparations illustrate two spatial dystrophin expression phenomena.**

Composite micrograph showing dystrophin immunostaining in a bulk preparation of EDL myofibers isolated from a single 60-week-old *mdx52-Xist*$^{\Delta hs}$ animal. Scale bars represent 1,000 µm, tiled images taken at 20× magnification and stitched together using LAS X. Nuclei were stained with DAPI. Both spatial dystrophin expression phenomena are apparent in this micrograph; (i) a 'zebra-like' banding pattern of patchy dystrophin expression, and (ii) the absence of dystrophin expression in centrally-nucleated myofibers and myofiber segments.

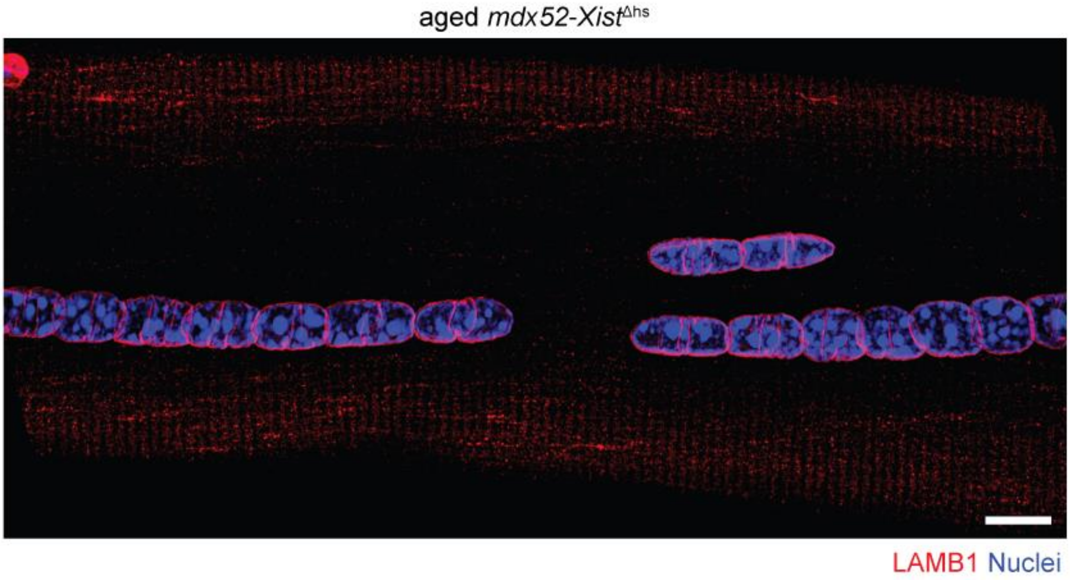


**Figure S5**

**Myonuclei in centrally-nucleated fibers are discrete and not fused.**

Representative micrographs of immunostaining for LAMB1 (lamin-B1) to show nuclear membrane organization in single isolated EDL myofibers from 60-week-old *mdx52-Xist*$^{\Delta hs}$ mice. The staining shows that the myonuclei exhibit discrete nuclear membranes suggesting that they are packed closely together, rather than being fused. Images taken at 40× magnification, scale bar represents 10 µm.

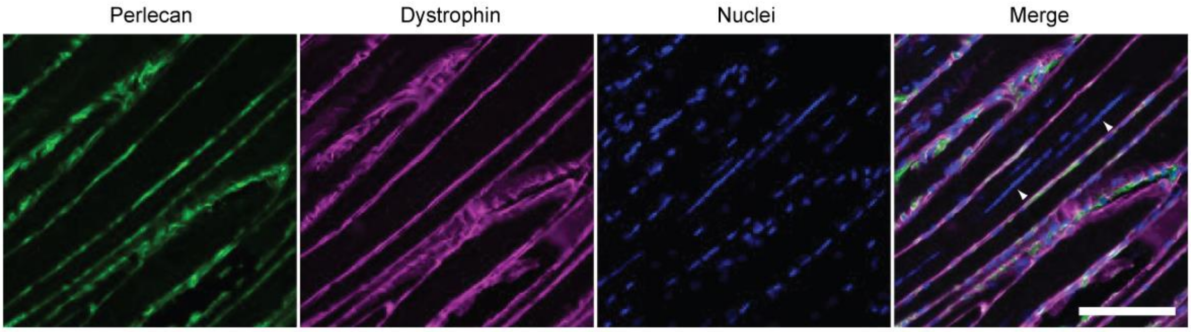


**Figure S6**

**Post-regeneration, centrally-nucleated myofibers express dystrophin in wild-type mice.**

Representative micrographs showing a longitudinal section through the tibialis anterior muscle of a $BaCl_2$-treated wild-type mouse (29 days post injury). The basement membrane was stained using antibodies against Perlecan, and nuclei were stained using Hoechst. Regions of centrally-located nuclei is highlighted with arrowheads. Images taken at 20× magnification, scale bar represents 100 µm.

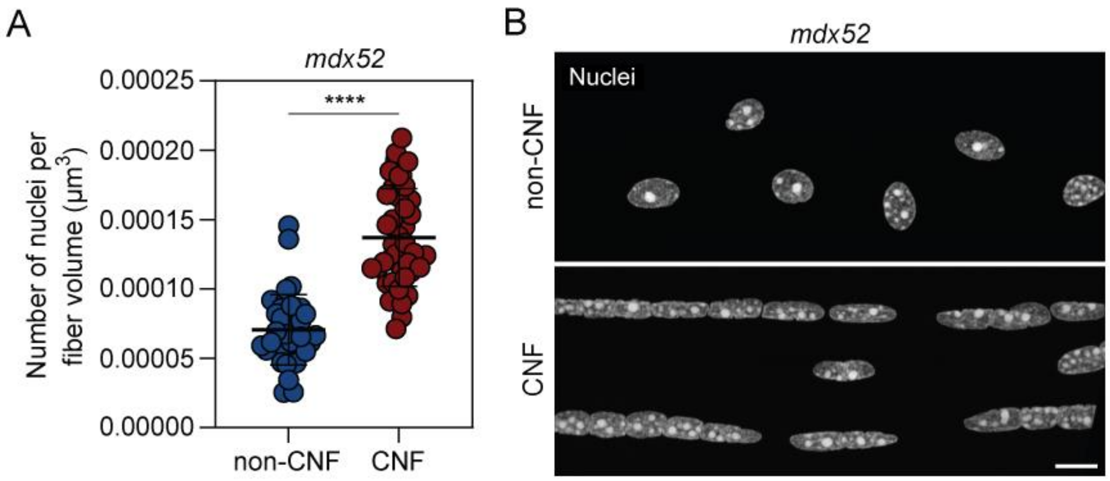


**Figure S7**

**Nuclei numbers are increased in *mdx52* centrally-nucleated myofiber segments.**

(**A**) Quantification of nuclei numbers per µm$^3$ myofiber volume in CNF (*n*=46) versus non-CNF (*n*=41) single isolated 12-week-old *mdx52* EDL myofibers. (**B**) Representative myonuclei staining of CNF vs. non-CNF myofiber segments. Images taken at ×40 magnification, scale bar represents 10 µm. Values are mean±SD. Statistical significance was assessed by Student's *t*-test, *****P*<0.0001.

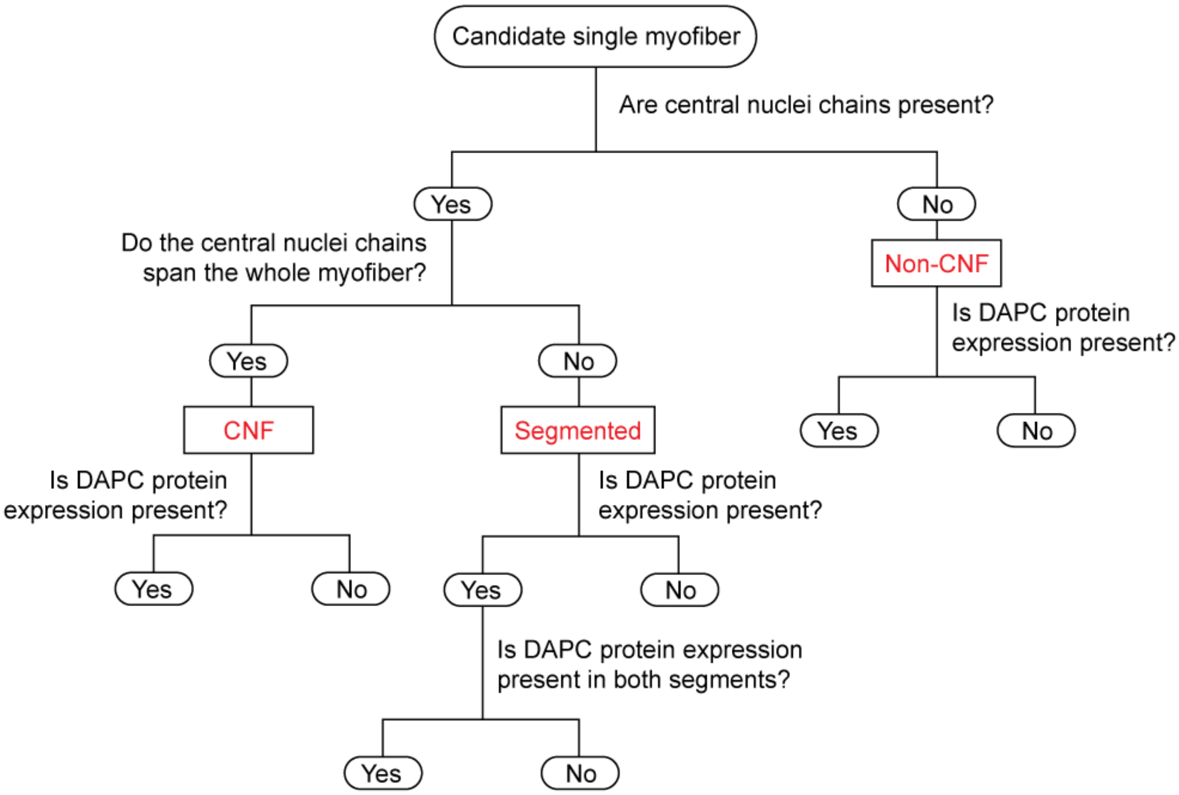


**Figure S8**

**Schema of single myofiber classification.**

Single isolated myofibers from *mdx52-Xist*$^{\Delta hs}$ mice were inspected and manually classified according to the above schema based on the degree of central nucleation and dystrophin/DAPC protein expression.

| Target | Host (clone) | Product ID | Manufacturer | Dilution |
|---|---|---|---|---|
| **Immunofluorescence** | | | | |
| **α-dystrobrevin** | rabbit pAb | α-1CTFP | In-house. Gift from Prof. K. E. Davies | 1:100 |
| **α-tubulin** | mouse mAb (AA13) | T8203 | Sigma-Aldrich | 1:1,000 |
| **α-tubulin** | rat mAb (YOL1/34) | ab6161 | Abcam | 1:250 |
| **α-tubulin** | rabbit mAb (EP1332Y) | ab52866 | Abcam | 1:250-1:500 |
| **Dystrophin (C-terminal)** | rabbit pAb | ab15277 | Abcam | 1:1,000 |
| **F-actin** | phalloidin probe conjugated with Alexa Fluor 568 | A12380 | Thermo Fisher Scientific | 1: 200,000 |
| **Laminin subunit α-2** | rat mAb (4H8-2) | L0663 | Sigma-Aldrich | 1:250 |
| **nNOS** | rabbit mAb | ab76067 | Abcam | 1:100 |
| **Telethonin** | rabbit mAb (EPR8375) | ab133646 | Abcam | 1:1,000 |
| **Titin** | mouse mAb (9D10) | 9 D10 | DSHB | 2-5 µg/ml |
| **Utrophin** | goat pAB | URD40 | In-house. Gift from Prof. K. E. Davies | 1:400-1:500 |
| **β-Dystroglycan** | mouse mAb (43DAG1/8D5) | NCL-b-DG | Leica Biosystems | 1:100 |
| **LAMB1** | Rabbit pAb | Ab16048 | Abcam | 1:200 |
| **Perlecan** | Rat mAb | A7L6 | Thermo Fisher Scientific | 1:1000 |
| **Western blot** | | | | |
| **Dystrophin (rod domain)** | mouse mAb (Dy4/6D3) | NCL-DYS1 | Leica Biosystems | 1:100 |
| **Vinculin** | mouse mAb (hVIN-1) | V9131 | Sigma-Aldrich | 1:100,000 |
| **Utrophin** | mouse mAb (8A4) | MANCHO3 | DSHB | 1:50 |

**Table S1**

**Primary antibodies used in this study.**

The anti-Utrophin antibody was obtained from the Developmental Studies Hybridoma Bank, created by the NICHD of the NIH and maintained at The University of Iowa, Department of Biology, Iowa City, IA 52242.

| Secondary antibody | Product ID | Manufacturer | Dilution |
|---|---|---|---|
| **Immunofluorescence** | | | |
| **Donkey anti-goat IgG Alexa Fluor 488** | A11055 | TFS | 1:500-1:1,000 |
| **Goat anti-mouse IgG Alexa Fluor-568** | A11004 | TFS | 1:500-1:1,000 |
| **Goat anti-mouse IgG Alexa Fluor-647** | A21235 | TFS | 1:500-1:1,000 |
| **Goat anti-mouse IgG Alexa Fluor-488** | A28175 | TFS | 1:500-1:1,000 |
| **Goat anti-rabbit IgG Alexa Fluor-594** | ab150080 | Abcam | 1:500-1:1,000 |
| **Goat anti-rabbit IgG Alexa Fluor-488** | A11008 | TFS | 1:500-1:1,000 |
| **Goat anti-rabbit IgG Alexa Fluor-568** | A11011 | TFS | 1:500-1:1,000 |
| **Goat anti-rat IgG Alexa Fluor-488** | ab150157 | Abcam | 1:500-1:1,000 |
| **Goat anti-rat IgG Alexa Fluor-647** | A21247 | TFS | 1:500-1:1,000 |
| **Goat anti-rat IgG Alexa Fluor-555** | A21434 | TFS | 1:500-1:1,000 |
| **Western blot** | | | |
| **Horse anti-mouse IgG-HRP** | 7076S | Cell Signalling | 1:10,000 |

**Table S2**

**Secondary antibodies used in this study.**

TFS, Thermo Fisher Scientific.

| Target | Product ID |
| --- | --- |
| mmu-miR-1a-3p | 002222 |
| mmu-miR-133a-3p | 002246 |
| mmu-miR-206-3p | 000510 |
| cel-miR-39 | 000200 |

**Table S3**

**List of Small RNA TaqMan assays used in this study.**

All assays were obtained from Thermo Fisher Scientific.
